# Investigating Clot-flow Interactions by Integrating Intravital Imaging with In Silico Modeling for Analysis of Flow, Transport, and Hemodynamic Forces

**DOI:** 10.1101/2023.06.03.543557

**Authors:** Chayut Teeraratkul, Maurizio Tomaiuolo, Timothy J. Stalker, Debanjan Mukherjee

## Abstract

As a blood clot forms, grows, deforms, and embolizes following a vascular injury, local clot-flow interactions lead to a highly dynamic flow environment. The local flow influences transport of biochemical species relevant for clotting, and determines the forces on the clot that in turn lead to clot deformation and embolization. Despite this central role, quantitative characterization of this dynamic clot-flow interaction and flow environment in the clot neighborhood remains a major challenge. Here, we propose an approach that integrates dynamic intravital imaging with computer geometric modeling and computational flow and transport modeling to develop a unified *in silico* framework to quantify the dynamic clot-flow interactions. We outline the development of the methodology referred to as Intravital Integrated In Silico Modeling or `IVISim`, and then demonstrate the method on a sample set of simulations comprising clot formation following laser injury in two mouse cremaster arteriole injury model data: one wild-type mouse case, and one diYF knockout mouse case. Simulation predictions are verified against experimental observations of transport of caged fluorescent Albumin (cAlb) in both models. Through these simulations, we illustrate how the `IVISim` methodology can provide insights into hemostatic processes, the role of flow and clot-flow interactions, and enable further investigations comparing and contrasting different biological model scenarios and parameter variations.

## 1 Introduction

Blood clot formation, growth, deformation, and embolization have an innate connection to the local flow environment [1]. Interactions of a continuously growing and deforming clot with surrounding pulsatile viscous blood flow at a vascular site *in vivo* leads to a complex dynamic flow environment, which determines flow-mediated transport in the clot neighborhood as well as flow-induced loading on the clot. Local flow environment plays a key role in determining transport of platelets and coagulation proteins, as well as transport of thrombolytic drug for treating stroke [2, 3]. Hemodynamic loading or forces such as pressure and shear on the clot can mediate clot growth, induce deformation, and lead to clot fragmentation and embolization [4, 5, 6]. Advanced understanding of these processes is essential for understanding disease etiology and progression for stroke and thrombosis, for evaluating efficacy of existing treatment approaches, and for evaluating novel treatment modalities including therapeutic targets for thrombolysis in stroke [7]. A wide-range of techniques have been employed to investigate the processes in clot formation. Experimental investigations include *in vivo* experiments in mouse models of stroke and injury [8, 9, 10, 11]; and microfluidic systems for coagulation including whole blood and platelet based assays [12, 13, 14]. Advancements in coupled multiscale multiphysics modeling techniques have enabled complementing these experimental approaches with high resolution computational models and *in silico* experimentation. Computational modeling has been successfully demonstrated for hemostasis and thrombosis phenomena across various investigations. These include: studies on developing highly resolved discrete mechanics models seeking to resolve detailed platelet-level interactions [15, 16, 17, 18]; studies on extending discrete platelet interactions with processes at larger scale to enable multi-scale analyses [19, 20]; and continuum averaged models for clotting phenomena including coagulation species transport and larger-scale clot-flow interactions [21, 22, 23, 24]. A further comprehensive review of state-of-the-art computational modeling is not the focus of this study, and the interested reader is referred to systematic reviews presented in [25, 26, 27]. Despite the range of investigations, quantitative understanding of unsteady flow and transport processes *in vivo* in the neighborhood of a dynamic clot of arbitrary shape and heterogeneous microstructure remains a major challenge. The combination of these factors is critical to resolve several aspects of clot structure-function relationship. Pulsatility, and arbitrary clot shape together can lead to complex flow structures around the clot owing to clot-flow interactions, which in turn can produce local flow features and barriers that organize mass transport [28, 29]. Clot structural heterogeneity such as pore-space distribution can determine the state of hindered diffusive transport of key species such as thrombin [30]. Formation of stable clots is necessary for effective hemostatic response. Conversely, for thrombosis, highly stable and stiff clots may be difficult to lyse, while low clot stability may increase chances of embolization [31]. Approaches to probe these features of clot structure-function relationship are therefore of high value.

Specifically, advancements in developing genetically modified mouse models, and imaging technology, have led to laser injury models in mice become a widely employed avenue to study hemostasis and thrombosis events *in vivo*. Typically, laser irradiation is used to create a localized vascular injury in mouse, and physiological clotting process *in vivo* is observed in real-time using techniques of intravital microscopy imaging [32]. Injury sites in existing studies have involved the mesenteric vessels, saphenous vein, and cremaster arterioles, with vessels in the range of 30-100 *μ*m. In a series of prior works [9, 17, 30, 33], laser mouse injury models have been systematically used to develop a systems level understanding of clot formation, growth, and biochemical transport and signaling in a coupled dynamic manner. In conjunction with simple computational analysis, clot geometry at different instants in time have been used to analyze clot deformation with the goal of identifying the transition point where large deformations and fragmentation initiates (referred to as the yield point, with the associated hemodynamic forces referred to as the yield stress) [34]. Combined with fluorescent tagging of platelets and key biochemical species, intravital mouse injury experiment data can provide a sufficiently detailed characterization of clotting phenomena immediately following the vascular injury. However, existing injury experiments do not directly include detailed unsteady flow information, and its coupling with associated biochemical transport dynamics and clot mechanics. This motivates the question: can we integrate intravital microscopy data with *in silico* modeling to quantitatively recreate the dynamic *in vivo* flow and transport processes around a growing and deforming clot? Here we address this question by developing an innovative computational multiphysics methodology that couples intravital imaging data with numerical flow and transport analysis named Intravital In Silico Integrated Modeling or `IVISim`. Our proposed methodology leverages a novel integration of an automated image-processing workflow with custom stabilized finite element algorithms to provide high resolution space-time varying flow and transport characterization around the dynamic clot post laser injury, that has been validated against experimental data. We demonstrate that through a framework like `IVISim`, we can directly generate a fully dynamic characterization of flow, transport, clot deformation, and flow-induced forces around the clot within a single unified computational toolkit, which is a significant advancement of the state-of-the-art in modeling thrombosis and hemostasis.

## 2 Methods

### 2.1 Intravital microscopy in a mouse laser injury model

We obtained time series images of hemostatic plug evolution from intravital microscopy of mouse laser injury experiments in mouse cremaster muscle arteriole [30]. Briefly, the experiments entailed using a laser to induce a perforation on the vessel lumen of live mice which triggered the hemostatic process. Confocal laser excitation was used to illuminate three fluorescent tracers: CD41, caged fluorescent albumin (cAlb), and P-Selectin, which are associated with hemostasis. CD41 signal intensity is associated with platelets that form the solid part of the clot. Region of high cAlb intensity indicates cell free plasma, including trapped plasma in the clot interstitial space or damaged vascular cells. P-selectin signal intensity indicates the region of highly activated platelets tightly packed near the perforation site. Consequently, high-intensity region of P-selectin indicates the “core” region of the clot [17]. For these experiments, 300 intravital image frames of each fluorescent channel were captured at 0.55 seconds per frame.

### 2.2 Automated segmentation workflow

The goal of this workflow was to identify and extract geometrically contiguous domain representing the region of high pixel intensity around the perforation site on each fluorescent channel. These were identified by a large continuous geometric envelope area of high pixel intensity value across each domain. We will refer to this domain as the “clot domain” for all three fluorescent channels. Due to the high number of image frames per stack, the segmentation method must be automated through the entire stack. The underlying steps within the overall workflow have been illustrated in Figure 1 panels a-d.

**Figure 1:**
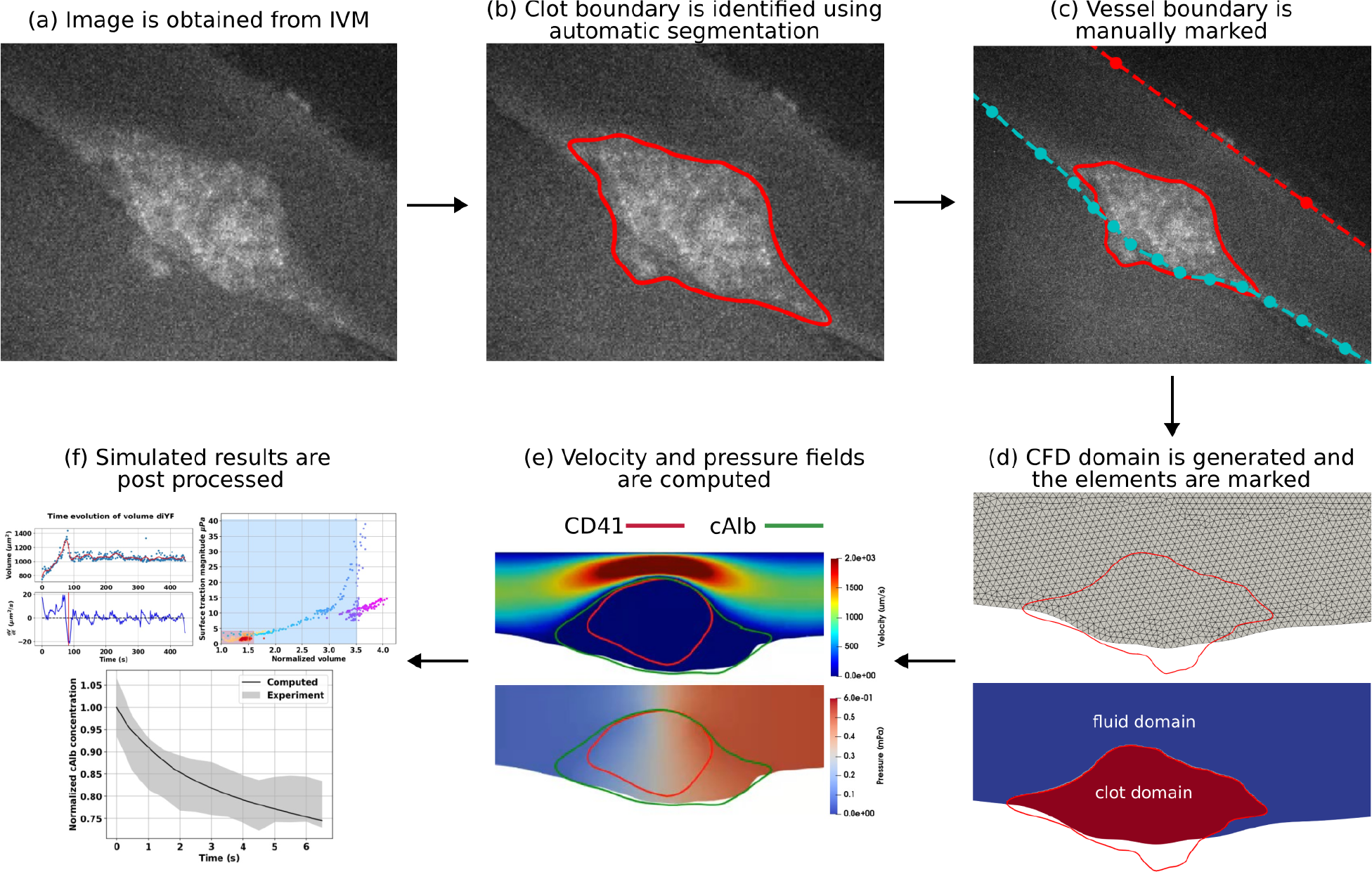
Schematic illustration of the `IVISim` workflow:(a) clot image is extracted from intravital microscopy images; (b) clot boundary is segmented at each frame of the image; (c) vessel boundary is marked from the images; (d) computational domain is generated, and points inside the boundary are marked; (e) hemodynamics around the clot domain is simulated; (f) simulated results are pre-processed to extract clot kinematics and mechanics.

#### 2.2.1 Image pre-processing

Each frame in the stack was pre-processed to remove any noise in the image. Since the pixel intensity value of the noise is often comparable to those inside the clot domain, applying a general-purpose denoising algorithm on the image directly does not remove the noise effectively. The image was first binarized based on the distribution of pixel intensity values. Binarizing the image differentiated the sparsely spaced bright pixels (noise) from the tightly packed bright pixels (clot) regions. The median filter was applied on the binarized version of the image to remove the sparsely spaced bright pixels. Finally, a Gaussian filter was applied to remove any additional noise and holes that may occur inside the clot domain. Image binarization and denoising were done using the open-source image analysis library OpenCV [35]. See Supplementary Material S1 for details on the binarization process.

#### 2.2.2 Image segmentation and clot boundary identification

The pre-processed image stack was segmented based on pixel intensity values using the k-means clustering algorithm implemented in the Python library Scikit-learn [36]. Since the image was already pre-processed, only two clusters were required to get the final clot domain segments. Finally, a Canny edge detection algorithm was applied to the segmented image to obtain the clot boundary pixels around the segmented portions. We observed that in some cases after segmentation, some image frames may contain multiple disconnected boundaries due to the remaining noise. To ensure that we get a single distinct clot domain in each frame, we selected the boundary with the largest perimeter as the final thrombus domain boundary. This process was efficiently automated by representing the image as a graph of boundary pixels. We then implemented a disconnected graph identification algorithm and use the largest sub-graph as the clot boundary across all of the stacks in the time-series of intravital images. See Supplementary Material S2 for additional details on the disconnected graph identification algorithm implemented in our framework. A simple analysis of quantitative accuracy of the segmentation output was conducted, by testing the segmentation workflow on a synthetic ellipsoidal patch geometry, with pixel image intensity statistics similar to the clot domain in the intravital image sequences as described in the Supplementary Material S6, and the errors of segmentation for varying ellipsoidal geometries was found to be ≤ 10%.

### 2.3 Reconstruction of clot dynamics

Pixel representation of the clot boundary is difficult to manipulate and is not directly suitable for a computational simulations due to the stair-step effect in the boundary model. A smooth curve is required to represent the physiologically realistic clot boundary. In `IVISim`, we parameterized the boundary as a B-spline curve representation using the `NURBS-Python` library [37]. We used the individual pixel positions as the control points of the curve. The time increments between each microscopy image frame obtained is 0.55 seconds. However, a smaller time increment was required to achieve a reasonably accurate flow and mass transport reconstruction over time. We obtained the clot boundary between each microscopy image frame by linearly interpolating the boundary points along the B-Spline curve which have the same angle from the center of mass of the clot region. An in-house ray tracing approach is used to identify these interpolating points on the B-Spline curve. The resulting smooth B-spline curve is independent of any background mesh used for subsequent finite element simulations, which is advantageous. See Supplementary Material S3 for implementation details.

### 2.4 Front tracking algorithm for moving clot domain marking

Since the shape of each clot domain are highly irregular and varies over time, efficient identification of finite element nodes inside the dynamic clot domains was a necessary step in developing the flow and transport simulations across multiple instances in time. Here, we adopted the point-set method as outlined in [38]. Briefly, this method involved solving Laplace’s equation ∇^2^*I* = 0 for an indicator function *I* on the fluid grid where the boundary conditions were such that *I* = 1 on each node which is part of an element in the mesh that contains a point on the segmented clot boundary. The nodes on the boundary of the domain were set to *I* = 0. Once this Laplace equation was solved, the nodal solution values were thresholded such that *I <* 1 − *ϵ* = 0 where *ϵ* is an arbitrarily small number. We note that this procedure adds one extra Laplace equation solve per time step in the entire image-processing workflow, which was a relatively cheap operation compared to the fluid flow and mass transport solutions as outlined in the later steps in the `IVISim` workflow. Furthermore, this front tracking step was only done once for each stack of microscopy images, rendering the associated computational expense to be small.

### 2.5 Moving fictitious domain stabilized finite element formulation

#### 2.5.1 Flow and transport modeling

Blood was assumed to be an incompressible, Newtonian fluid with a constant averaged dynamic viscosity of *μ*= 4.0 cP, and a constant average density of *ρ* = 1.06 g/cc. Flow around the clot was computed using the unsteady Navier-Stokes equation. Flow inside the heterogeneous porous clot domain was modeled using the Brinkman equation for low Reynolds number flow through porous media [39]. The overall flow and biochemical species transport using the Brinkman Navier-Stokes formulation was numerically simulated based on a custom stabilized finite element algorithm adopting techniques from prior works [40, 41]. Specifically, the final system of mass and momentum balance equations was solved using a Petrov-Galerkin stabilized finite element method. For modeling the Brinkman resistance to the flow, the clot permeability was estimated as a function of porosity using the classical form of the Kozeny-Carman relation stated as follows:

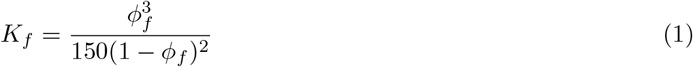

where *K*_*f*_ is the permeability estimate, and *ϕ*_*f*_ is the averaged porosity in a region of the porous clot domain being considered. We note that this relationship is only an approximation and may not be suitable for all clot scenarios. Specifically, in prior work [42] it has been reported that platelet rich clot permeability is better approximated by direct Kozeny-Carman type relations, while fibrin composition may influence these permeability estimates and require additional terms. These have not been discussed in our current study, but are easy extensions of the existing `IVISim` framework requiring simple algebraic modifications of the permeability-porosity relationship. The computed flow velocity within and around the dynamic clot was used as a background flow for numerically simulating unsteady biochemical species transport for the clot domain using an advection-diffusion formulation. Within `IVISim`, the resulting mass transport equations are solved using a custom finite element algorithm that combines Petrov-Galerkin stabilization with concentration jump stabilization techniques. This custom stabilization algorithm enables `IVISim` to handle more complex spatiotemporally varying porosity and diffusivity inputs, if such data is available for the dynamic clot domain. Details of the stabilized finite element algorithms and their mathematical formulations are included in the Supplementary Material S4. The combined stabilized finite element algorithms were implemented using an open-source finite element library `FEniCS` [43], with Python as the primary scripting language.

#### 2.5.2 Dynamic clot domain properties

Using the overall image-processing framework in `IVISim` we reconstruct the clot as a dynamic heterogeneous porous domain. This enables a simplified averaged form of capturing clot microstructural features that vary in time. Essentially, this implies that for our numerical formulation, the porosity and diffusion coefficients are actually represented as: *ϕ* (***<u>x</u>***, *t*), and *D* (***<u>x</u>***, *t*). We note that in this study, we have not attempted to quantitatively describe the fully-resolved spatio-temporally varying clot pore network and microstructure information, and obtain mathematically detailed definitions of these functions. However, when such information is available, the corresponding *D* and *ϕ* functions can be used as inputs to the framework outlined here. For the cases and analyses presented here, we represent the clot domain varying in time as Ω^*T*^ (*t*). We have dynamic reconstructions enabled from the CD41 signal intensity which gives us the overall clot domain, and the P-selectin signal intensity which gives us the core region of the clot. Representing the dynamic clot core domain as: 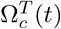, we obtain the dynamic shell domain as: 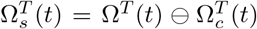. The porosity is then defined as an implicit function of space and time through the domain information as follows:

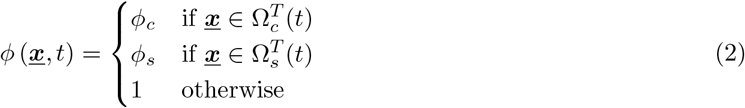

Likewise, the diffusivity values are also specified in form of effective diffusivity values in the clot interior. This accounts for observations in previous studies that reported hindered diffusion to be the dominant mode of mass transport inside the clot [17].

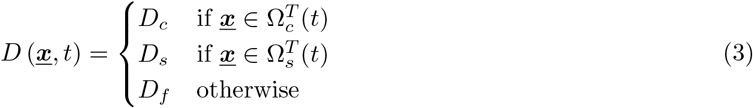

where *D*_*f*_ is the free-stream diffusion coefficient of the biochemical species in consideration (for example, cAlb in this study, as shown later), and *D*_*c*_, *D*_*s*_ are core and shell diffusion coefficient that are scaled versions of the free-stream diffusion coefficient using scaling relations based on porosity and pore-connections as available from theories of transport in heterogeneous materials [44, 45, 46]: for example, for the core: *D*_*c*_ = *α*(*ϕ*_*c*_(***x***, *t*))*D*_*f*_ with *α* being a porosity dependent scaling factor; and likewise for the shell.

### 2.6 Simulation case-studies

Here we demonstrate our complete `IVISim` methodology as illustrated in Figure 1, using a simulation casestudy involving intravital data from two mouse models reported in prior work [30]. The first set was standard wild type (WT) male mice 8-12 weeks of age prepared as outlined in [30, 17]. The second set involved mice with a substitution of phenylalanine for 2 tyrosine residues in the *α*_*IIb*_*β*_3_ integrin, referred to as diYF knockout mice. One representative set of intravital microscopy data for each mouse strain was used for this study. This combination of a WT and a diYF knockout model was chosen here to demonstrate the `IVISim` predictions on two clots with significantly different structural features and dynamics. The diYF knockout is known to have impaired integrin outside-in signaling and hindered *in vitro* clot contraction. For both, caged fluorescein conjugated to Albumin (cAlb) was imaged, to quantify transport rates in the clot domain, as described in [9]. For each model, the cremaster arteriole centerline flow velocity was estimated experimentally from the images to be 1300 *μm/s*, measured by tracking tagged platelets in the vessel before the laser injury was made. For simulations, this flow was specified as an inflow boundary condition mapped on to the inlet face of the arteriole domain as a parabolic flow profile. Fixed averaged viscosity and density values of 4.0 *cP* and 1.06 *g/cc* respectively were assumed. Porosity values in clot interior were taken to be in the range 0.2-0.4 as reported in [17]. Specifically, porosities in the dynamically evolving core and shell regions for the WT model were assigned to be 0.2 and 0.3 respectively, while those for the diYF model were assigned to be 0.2 and 0.4 respectively. For cAlb mass transport simulation, we assumed a diffusion coefficient of 60 *μm*^2^*/s* in the free flow domain as also reported in prior work [17]. Inside the thrombus domain, mass transport is hindered due to the thrombus microstructure as detailed in several previous studies. Hence, the effective diffusion coefficient was modeled as a scaled version of the corresponding free-stream value. For this study, the scaling comprised of a porosity and pore-tortuosity based factor as outlined Section 2.5, as well as an additional geometric correction factor that accounted for the comparison between the exact reconstructed geometries of the clot and the ellipsoidal approximated models reported in prior work [17]. The resultant effective diffusion coefficients for the clot interior used in this simulation were 0.8 *μm*^2^*/s* in the shell region and 0.4 *μm*^2^*/s* in the core region for the WT variant; and 1.0 *μm*^2^*/s* in the shell region and 0.5 *μm*^2^*/s* in the core region for the diYF variant. We note that these effective diffusion coefficients are different from the values reported in the prior work [17] due to the mismatch in the exact vs ellipsoidal geometry. To enable further analysis of parameter sensitivity and uncertainty quantification, we implemented a workflow to integrate `IVISim` with the open source toolkit `Dakota` [47]. We conducted an analysis of sensitivity of the segmented clot domain with respect to key image processing parameters using `Dakota`’s built-in Latin Hypercube Sampling approach. Details of this methodology is outlined in the Supplementary Material S6. Finally, to demonstrate the quantitative performance of the dynamic clot interpolation workflow within `IVISim`, we down-sampled the dynamic image set by removing every two frames from the image stack, reinterpolated the temporally down-sampled images between each frame, and compared the output against the clot boundary obtained from the original image stack. Additional details on this methodology are outlined in the Supplementary Material S3.

## 3 Results

### 3.1 Characterization of *in vivo* dynamic flow environment around a clot

Our first objective was to demonstrate that the resultant `IVISim` framework enables a quantitative characterization of the *in vivo* dynamic flow environment around a growing and deforming clot, which is otherwise not available from the intravital microscopy alone. To establish this, we used our *in silico* framework to compute the space-time varying flow around the hemostatic plug formed at the puncture injury site in the mouse cremaster arteriole. The resultant flow and clot shape data for the WT mouse are presented in Figure 2. Specifically, panel a. illustrates the spatio-temporally varying hemodynamics in the clot neighborhood. We observe that during the early stages of the clot formation, where platelet aggregation is still nascent, and the core-shell structure of the clot is evolving, the flow remains mostly undisturbed, and permeates the clot interior. As the clot grows, and organizes into a distinct structure comprising a tightly packed core and a loosely packed shell, the flow is distinguishable into two regimes. There is a significantly arrested, slow intrathrombus flow in the heterogeneous pore space of the core and the shell of the clot, with the core being less porous, and therefore showing more arrested flow than the shell region. External to the clot, the resistance to the flow offered by the clot as it grows and occludes greater portion of the vessel lumen leads to rapid increase in flow velocity at the regions of high occlusion. These observations are consistent with multiple prior studies on clot-hemodynamics interactions. The overall Reynolds number associated with the computed flow environment was in the range of ≈ 10^−2^-10^−3^, consistent with numbers reported in prior work [17]. Additionally, we observe a distinct region of slow pooled flow with high cAlb signals, that originates as the injury itself leads to a local deformation of the vessel lumen, and subsequently a stagnation region for the flow.

**Figure 2:**
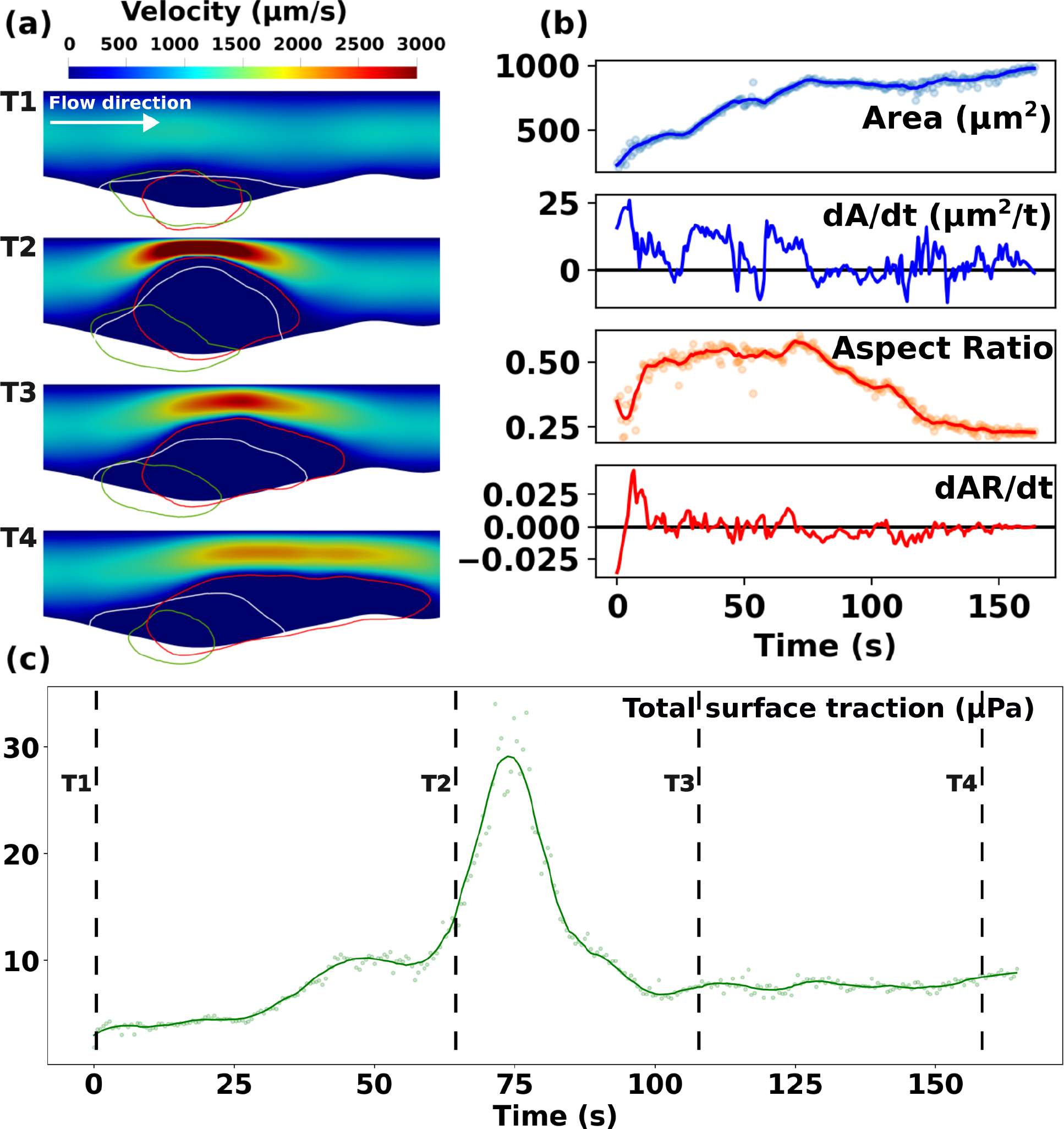
Flow and clot kinematics data obtained from `IVISim` simulations on wild-type (WT) mouse laser injury data. Panel a. shows the unsteady flow environment at 4 consecutive instances in time (T1-T4). Red curve depicts the shell, Green depicts the core, and White depicts the plasma cAlb domain. Panel b. depicts the clot shape variations over time tracked using clot domain area and aspect ratio. Panel c. presents the total flow-mediated traction forces (shear and pressure) along the clot boundary over time.

In addition, a continuous sequence of clot geometries is extracted using our automated algorithm, which recreates the continuous kinematics of the clot through the intravital image acquisition time, details of which are presented in Figure 2, panel b. for the WT case. The tracked clot kinematics is used to compute clot shape and size features. We note that within `IVISim`, high frequency noise content due to imaging artifacts are filtered out to ensure that only contributors to clot kinematics can be observed. The reconstructed dynamic clot domain is robust to the choice of the key image processing parameters, and the sensitivity analysis via integration with `Dakota` as described in Section 2.6 shows that the average variabilities in reconstructed area and aspect ratio are less than ≈ 10% and ≈ 2% respectively (see also Supplementary Material Figure S11. Additionally, comparisons of the interpolated boundary from the image-stack with those obtained from a down-sampled sequence as described in Section 2.6 demonstrated that the interpolation algorithm was robust. The point-wise averaged distance between the down-sampled interpolated boundary and the original boundary is in the order of *<* 1.0 microns for both WT and diYF models (see Supplementary Material Figures S4 and S5). The unsteady kinematics is a result of clot growth and deformation under flow, as well as clot contraction, and fragmentation or embolization due to flow-induced loading. We note that these occur at differing time-scales, with events like growth and stretching/deformation occurring over a slower time scale compared to embolization. Additionally, embolization in particular is a discrete phenomenon associated with a rapid change in clot volume. Motivated by these arguments, we also computed the rate of change of clot shape and size, tracking not only the magnitudes but also positive/negative rates of change as indicators of clot kinematics. The computed trends in overall area are the same as those reported in prior works [30, 34] which do not use our automated technique, hence validating our geometric/kinematic reconstruction approach. Specifically, we observe that the overall clot domain (represented by the CD41 channel signals in these datasets) reproduces the characteristic growth behavior followed by an inflection in the geometry and a slower growth or a plateau phase as observed in experiments in [30]. We note that the resulting clot kinematics reported in [30] utilized a pixel intensity thresholding based technique, while our approach seeks to identify a coherent geometric envelope or domain of the clot. Consequently, there could be regions inside the actual core/shell domain that are not illuminated in a pixel intensity threshold image, but will still be resolved using our approach owing to the pixel clustering approach. Additionally, in our methodology we obtain the overall clot domain with pores, whereas if we purely threshold based on pixel signal intensity, we are not guaranteed to obtain the manifold with pore spaces inside. Hence, in summary, between the approach and final kinematics reported in prior work [30], and those represented in our work, we are capturing the overall domain geometry differently. While acquiring and comparing the averaged trends across multiple experiments for a direct quantitative comparison of the clot kinematics was not the goal here, we have included a sample quantitative comparison against WT CD41 dataset trends in Supplementary Material in Section S5. The factors identified above pertaining to differences in methodologies account for the differences observed between the datasets, while still demonstrating the similarities in the temporal trends as reported earlier in this paragraph.

Furthermore, computed flow data was post-processed to obtain the flow-induced loading on the clot. Specifically, in Figure 2, panel c. we illustrate the total flow-induced loading on the clot generated by clot-flow interactions over time. In accordance with fluid-structure interaction principles, we compute the total force on the clot as the integral of the total stress on the clot boundary originating from clot-flow interactions. This includes the total pressure and total shear loading obtained by integrating pressure and shear from the flow along the boundary of the clot over time. The total flow-induced loading is a key driver of clot structural deformation and embolization. Additionally, as demonstrated in our prior works [48, 28, 29], the pressure gradient across the clot boundary is a determinant of the extent of permeation of biochemical species into and out of the clot domain at the permeable shell boundary of the clot. Specifically, for the WT model, we observe that the loading increases, with increase in occlusion, until a peak value at which point the clot deforms, relieving the load. Comparing with the kinematics data in panel b., we note that this point of peak loading corresponds to instances where the clot area over time slows down, and the aspect ratio reduces, indicating geometrically that the WT clot essentially deformed and stretched under the action of the hemodynamic load, but not necessarily embolized. This observation enables a distinction between deformation and embolization of a clot *in vivo*, which we address further in Section 3.3 and 3.4.

### 3.2 Validation of model predictions using cAlb transport

Another key objective in this study was to demonstrate the validity of the predictions from the complete `IVISim` numerical framework. For this purpose, we implemented an *in silico* model for transport within and around the clot tracked by cAlb, and compared the simulated cAlb concentration variation against cAlb concentrations measured *in vivo* using fluorescent microscopy. Specifically, as described in Section 2.5, for the intravital images periodic pulses of 405-nm light were used to uncage cAlb and the resulting fluorescence intensity and decay in the clot domain were monitored with fluorescent microscopy. For the *in silico* model, cAlb concentration was initialized over the complete clot domain, and the entire `IVISim` workflow was executed to obtain a simulated cAlb distribution varying in space and time. Figure 3 panel a. illustrates the computed distribution of cAlb at varying instants in time between two successive pulses. The computed distribution reveals an initial washout of cAlb concentration in the vessel lumen, followed by a slow variation of concentration driven by diffusion dominant transport of cAlb in the clot interior and flow-mediated permeation of cAlb from the clot interior into the lumen. This leads to an overall decay curve for the cAlb concentration as presented in Figure 3, panel b., which has been reported in prior works [9]. The resultant cAlb concentration decay was compared with the decay behavior obtained from intravital imaging, and the predictions from `IVISim` framework agree well with the expected averaged decay behavior observed across multiple experimental runs for the WT mouse cremaster injury as shown in panel b. The hemostatic clot manifests significant heterogeneity in micro-composition, which is reflected in the cAlb concentration map as well. It is observed that majority of the permeation-driven change in cAlb concentration is in the shell region of the clot, while within the core the cAlb remains arrested with much slower diffusive transport leading to lower rates of cAlb concentration decay.

**Figure 3:**
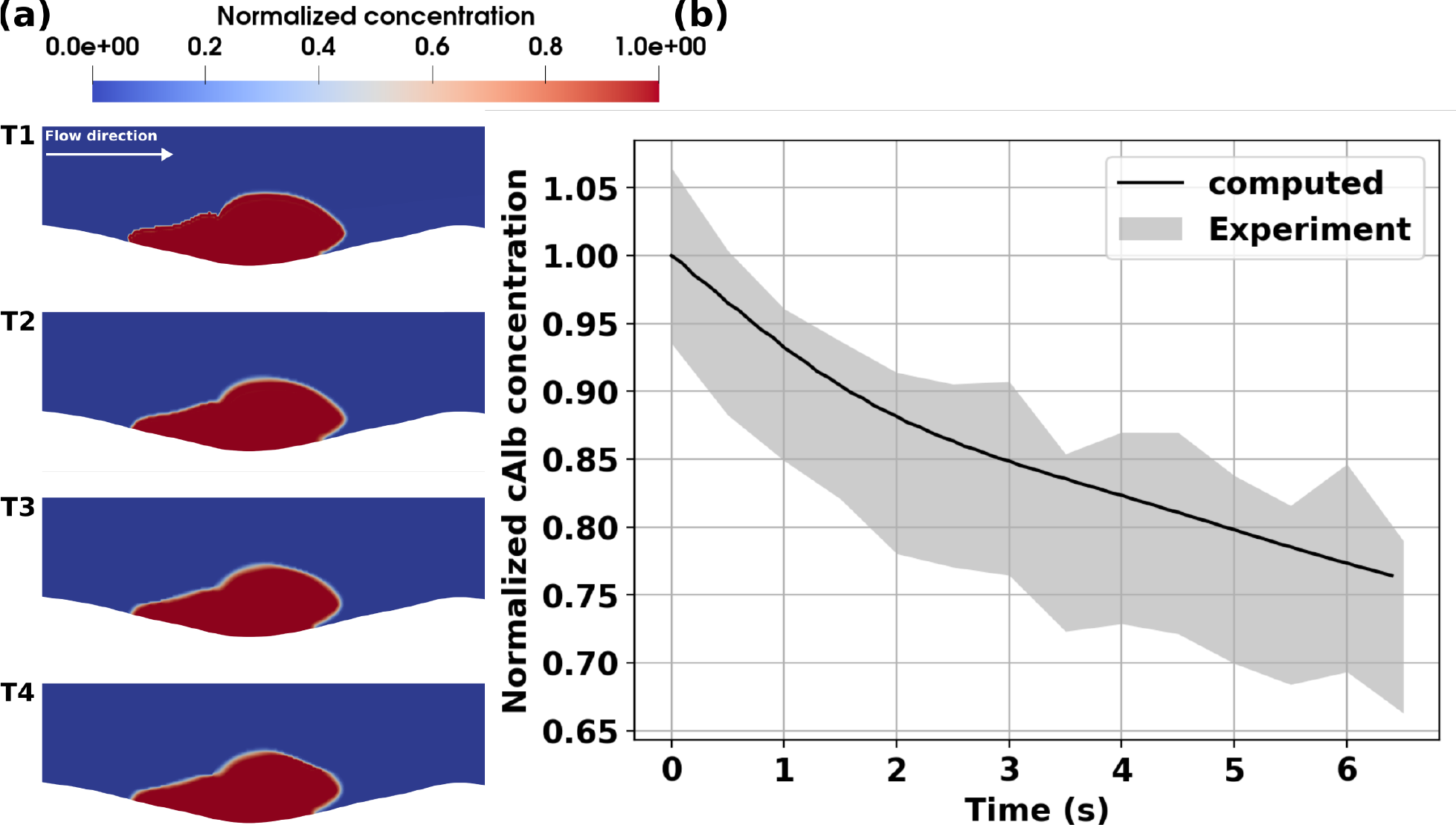
Simulation of cAlb for the wild-type (WT) mouse injury model. Panel a. depicts predictions from `IVISim` for spatial distribution of cAlb across 4 consecutive instants in time (T1-T4). Panel b. presents a volume averaged decay of cAlb within the clot domain, which is compared against experimentally measured and quantified decay behavior. The agreement between predicted curve and the experimental range helps validate `IVISim` predictions for cAlb transport.

### 3.3 Comparing clot-flow interactions across the WT and diYF models

In Figure 4, we present the computed flow around the hemostatic clot in a diYF knockout mouse, with the objective of comparing the flow and transport environment between the WT and diYF knockout models. Similar to Figure 2: (1) panel a. presents the space-time varying hemodynamic patterns around the clot; (2) panel b. depicts the time history of clot shape and size, and their rates of change, obtained using `IVISim` automated algorithm; and (3) panel c. illustrates the flow-induced shear and pressure loading integrated along the boundary of the growing clot over time. When compared to the hemodynamics in the WT clot neighborhood, few qualitative similarities were noted, such as: (a) faster flow around the clot as it grows; (b) slower arrested flow in the clot interior; and (c) a region of pooled stagnated flow at the base of the clot formed due to the vessel deformation at injury site. However, the clot in the diYF knockout model grows to a much smaller size before starting to embolize, as indicated by the drop in clot area at the instant of peak hemodynamic loading, with minimal change in clot aspect ratio. Subsequently, the clot sheds emboli continuously under the action of the flow-induced loading (*see also animations provided in Supplementary Material S7*). Consequently, the peak hemodynamic loading is substantially lower in the diYF knockout model than for the WT case. On the other hand in the WT mouse considered in our simulations, the hemostatic clot is able to grow to a bigger structure that can sustain a higher level of flow-induced loading.

**Figure 4:**
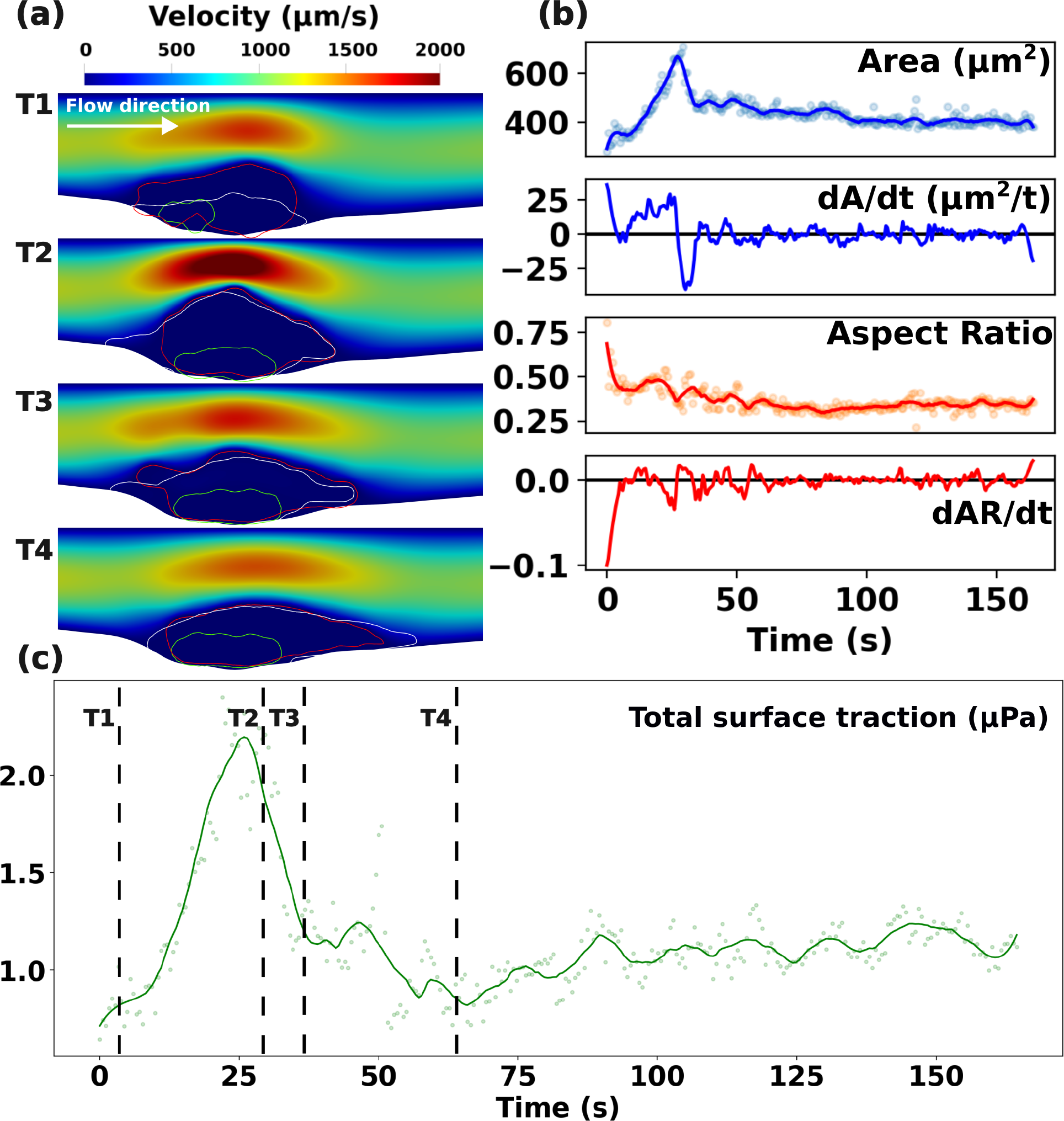
Flow and clot kinematics data obtained from `IVISim` simulations on a diYF knockout mouse laser injury data. Panel a. shows the unsteady flow environment at 4 consecutive instances in time (T1-T4). Red curve depicts the shell, Green depicts the core, and White depicts the plasma cAlb domain. Panel b. depicts the clot shape variations over time tracked using clot domain area and aspect ratio. Panel c. presents the total flow-mediated traction forces (shear and pressure) along the clot boundary over time.

This can be further illustrated by looking at the time evolution of the regions of CD-41 and P-selectin activity as presented in Figure 5. For the WT simulations shown in panel a., the region with high P-selectin intensity indicating the core of the clot, grows and shrinks continuously, with a significant reduction in area being observed after the initial period of growth. The region of CD-41 activity continues to grow, and deform, but no reduction in area is observed. Conversely, for the diYF knockout simulations shown in panel b., the P-selectin region shows minimal levels of area reduction, and the CD-41 region shows embolization leading to a sharp drop in volume, as opposed to more flow-induced deformation which is stably handled by the WT clot. We note that the diYF knockout mice are expected to have deficient clot retraction *in vivo*, and the above observations on clot growth, embolization, and growth/shrinkage of P-selection vs CD-41 intensity envelopes can be related to differences in clot retraction behavior. Potentially the deficient retraction in diYF knockout cases may also alter the clot structural integrity, as discussed in several studies investigating clot retraction and its role [49]. However, while the simulations presented here indicate that the `IVISIm` methodology is capable of detecting such structure-function differences in the hemostatic clot, additional computational experimentation with a larger set of mouse samples will be required to infer generalized conclusions about these aspects of clot structure-function relationship.

**Figure 5:**
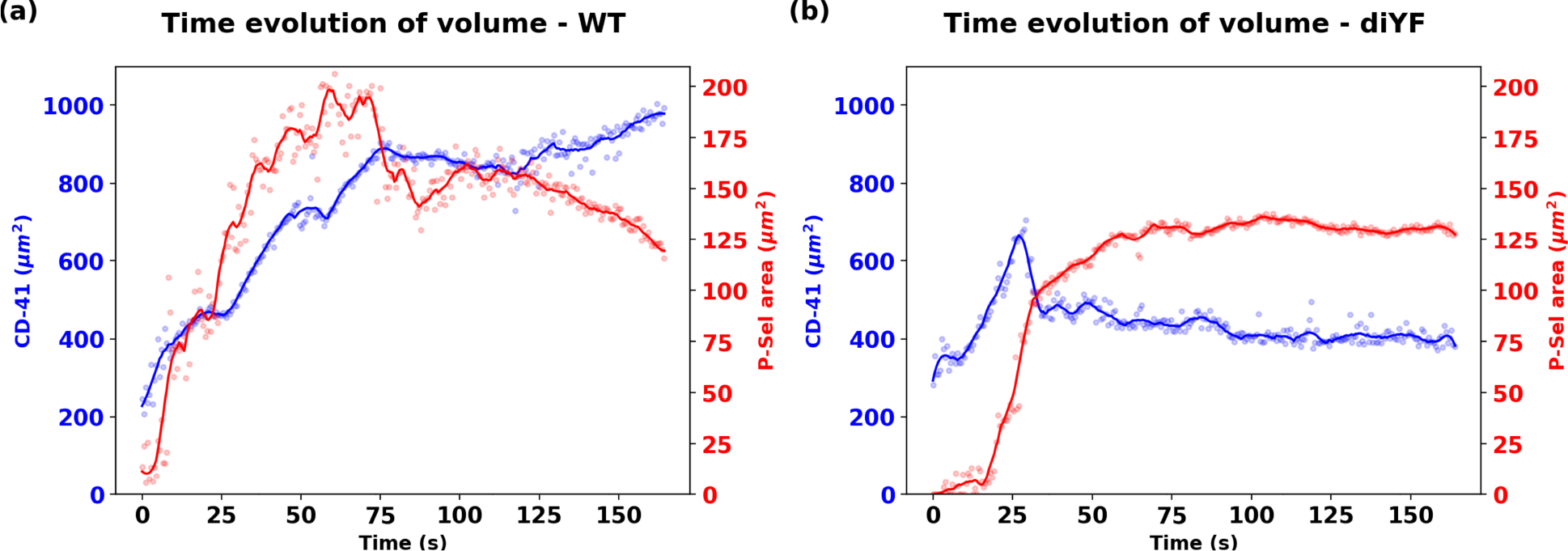
Comparison between the overall clot domain (denoted by CD41 channel image intensity) and the core of the clot (denoted by P-Selectin channel image intensity) between the wild-type and the diYF knockout mouse model clots.

Additionally, for the diYF knockout case, the distribution of cAlb in the clot domain, and the decay of averaged intra-clot cAlb concentration are presented in Figure 6, panels a. and b. respectively. The distribution shows slow variations in concentration over the small core region, but significantly rapid variations over the shell region of the clot, when compared to the WT case. This is supported by a faster decay rate of the averaged cAlb concentration compared to WT case as indicated in Figures 3 and 6 panel b. In both cases, the core region with higher extent of compaction traps the species concentration, while the shell region manifests the greater extent of transport and cAlb species washout. Overall, the simulations indicate that with impaired contraction, and associated smaller core volume and overall higher porosity, the solute transport rates in the diYF knockout cases can significantly differ as compared to WT mice [30, 17].

**Figure 6:**
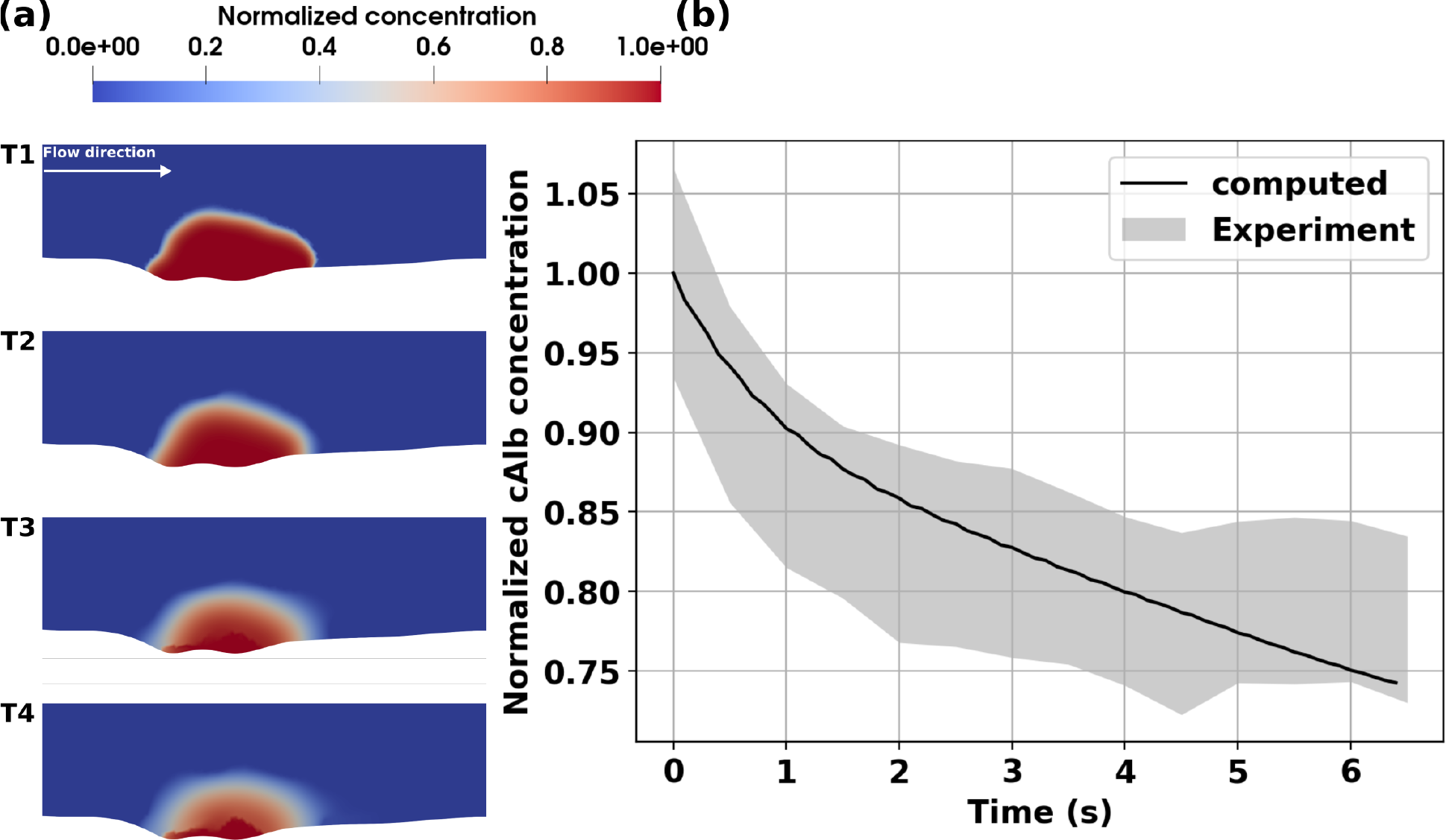
Simulation of cAlb for the diYF knockout mouse injury model. Panel a. depicts predictions from `IVISim` for spatial distribution of cAlb across 4 consecutive instants in time (T1-T4). Panel b. presents a volume averaged decay of cAlb within the clot domain, which is compared against experimentally measured and quantified decay behavior. The agreement between predicted curve and the experimental range helps validate `IVISim` predictions for cAlb transport.

### 3.4 Characterization of flow-induced clot deformation properties

Here, we attempt to summarize the key quantifications from the `IVISim` framework into a visual tool for characterization of clot-flow interactions *in vivo*. For this, we plot the continuous time-history of clot size against the continuous time history of flow-induced loading integrated along the dynamic clot boundary, for each instant in time. This is a form of mapping of a force-volume relationship *in vivo* as predicted by the `IVISim` framework, leading to a dynamic force-flow-volume phase-space representation of the clotting process. The resultant plot is shown in Figure 7, where the mapping of total flow-induced loading (that is, combined pressure and shear loading) for both WT and diYF knockout cases are overlaid together. See also Supplementary Data S5 and S7 for additional visuals illustrating the pressure and shear loading individually. The point of embolization is identified (guided by the time history plots as shown in Figures 2 and 4 earlier) as the point of inflection in the clot size as function of time, indicated using an arrow for both WT and diYF models in Figure 7. The corresponding hemodynamic loading that initiates embolization can be directly estimated as the intercept along the horizontal axis at this point of embolization. Furthermore, a bounding box is marked out between the point of clot initiation, and the point of embolization, represented as the shaded rectangular regions in the Figure 7. Comparison between the WT and diYF knockout mouse models considered in this study show that the WT hemostatic clot grows to a bigger size and sustained a higher loading prior to embolization, causing this bounding box to be bigger along both x and y-axis as compared to that for the diYF knockout case. While this is true for both the shear and pressure forces on the clot, the differences are more pronounced when the pressure forces are considered. This is likely due to the fact that these forces are computed as an integral over the entire clot boundary, and while shear is higher at the regions of peak occlusion along the boundary (affecting the forces only locally), pressure will be determined more by the resistance offered by the clot to the flow (affecting the forces more widely along the clot boundary).

**Figure 7:**
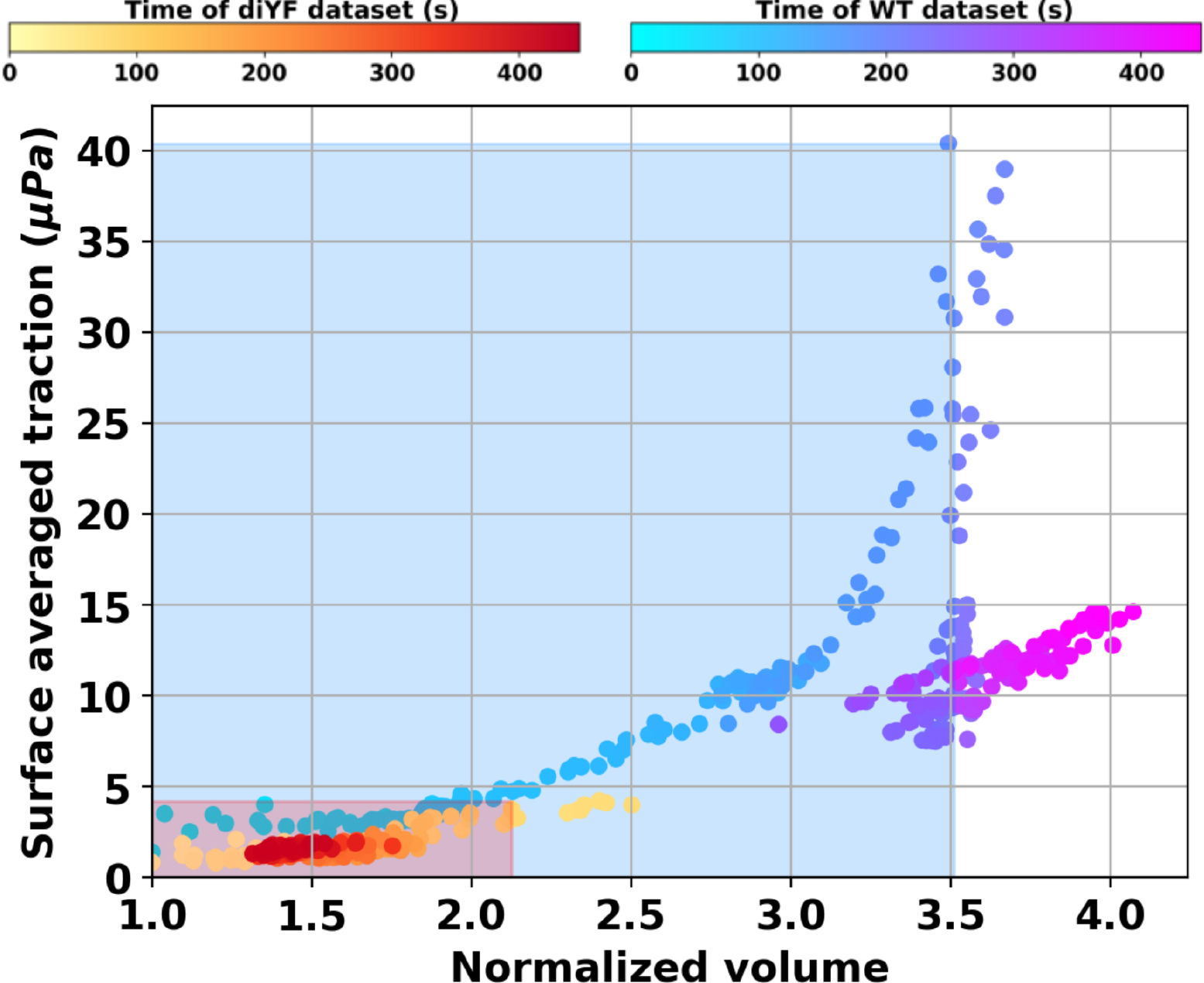
Collapsing the force-deformation behavior of the clots formed in the wild-type and the diYF knockout mouse injury models. Each point in the plot represents the flow-mediated force and corresponding clot size at an instance in time. Dynamic tracking of these quantities leads to a phase-space plot in the hyperspace formed from force magnitude and clot volume. The bounding boxes represent the force and size at the point of embolization in each case, which is indicated as the upper right corner of the bounding box for each model marked with an arrow.

## 4 Discussion

Here we have outlined the development of the `IVISim` framework, which is capable of directly integrating flow and transport physics models with dynamic intravital microscopy. Through a set of simulation case studies, we demonstrated that this approach is able to provide detailed quantitative flow and transport information for the *in vivo* murine thrombus neighborhood. This flow environment information is not directly available from microscopy alone. For models based on 2D intravital image sequences, the entire `IVISim` computational workflow can run on standard desktop computing resources, making this a valuable utility complementing standard intravital experiments. Real *in vivo* clotting process involves a complex, dynamic, and synergistic coupling of clot structure, clot-flow interactions, flow-mediated transport, and biochemical processes. The integration of intravital imaging and detailed quantitative *in silico* modeling enables us to potentially investigate these dynamically coupled set of phenomena. This has remained a major challenge, and our study indicates that a data-driven *in silico* approach like `IVISim` can substantially extend the insights that intravital imaging can provide. Additionally, the methodology developed here is not limited to mouse intravital microscopy alone, but also generalizable to dynamic imaging of clotting processes as obtained from microfluidic human whole blood assays, thereby making it a broadly applicable tool. Furthermore, the insights from murine *in vivo* and human whole blood *in vitro* clotting investigations can be integrated by the simulation capabilities embedded in `IVISim` framework, to explore a variety of *‘what if ‘* scenarios for the human *in vivo* clot environment, and flow and transport phenomena therein. Such quantitative information on human clot environment *in vivo* is otherwise significantly challenging to obtain, and motivate future investigations on using `IVISim` for *in silico* exploration of human clotting phenomena. One relevant limitation to note here is that we have demonstrated our methodology on 2D models of the clot as opposed to actual 3D models. This is not actually a limitation inherent in the `IVISim` framework, but rather is attributed to the type of intravital data available. When 3D data are available, the automated kinematic reconstruction processes can be extended by incorporating fully 3D point cloud morphing techniques such as coherent point drift [50].

Additionally, we demonstrate that using this framework we can computationally estimate the hemodynamic loading generated from the clot-flow interactions. In our simulations, we investigate the trends in the total hemodynamic loading which is a combination of the pressure and shear loading, instead of considering them in isolation. We observed the loading generated by pressure forces to be substantially greater compared to that generated by shear. Flow-induced shear loading on the clot is high locally at the location of maximum occlusion. However, pressure increases more globally as the occlusive clot resists flow, and drives up the pressure differential required to push the fluid across the occlusion region. One simulation parameter that can significantly influence the force magnitudes is estimates of vessel flow rates or flow velocity. Here we have used a peak centerline velocity of 1300 *μm/sec* as obtained from tracking of tagged platelets in the vessel prior to laser injury [17]. This velocity can be different depending upon the flow conditions post-injury, which may alter the specific numerical values of the forces, although we expect the trends reported in Figure 7 to remain similar since the clot deformation kinematics data is directly computed from the image. In this context, we note that in real vascular networks, in the presence of branching vessels, the above flow-induced loading behavior may be different, as the flow may reroute through the branching vessels and assume a path of least resistance. Such vessel branching effects have not been incorporated in the analysis presented here, but comprise an important feature of the *in vivo* clot environment, and needs to be accounted for in future investigations. Quantifying the flow-induced total loading, instead of pressure or shear alone, is therefore advantageous in encapsulating such variability.

Specifically with the laser puncture injury models considered here, for both simulation cases we observed the presence of a prominent region of slow arrested flow at the base of the clot near the vessel wall injury site, with high cAlb fluorescence signal intensity outside of the platelet clot domain (that is, regions indicated by the P-selectin and CD41 signals). This region serves partly as an extension to the shell region of the clot (and consequently referred to as an ***extended shell***) as indicated in a sample illustration from the diYF clot model in Figure 8. The left panel illustrates the formation of this extended shell as identified through the images (*marked using the label (a)*), and the right panel establishes this region with slow flow velocities (*marked using the label (b)*). This slow stagnated flow region originates from the vessel deformation *in vivo* at the injury site, and the flow resistance from the hemostatic plug that forms around the injury site. Our simulations indicate that: (a) this extended shell varies with clot growth and deformation; (b) can influence biochemical species transport in and out of the clot environment; and (c) while not directly a part of the clot itself, it potentially comprises fibrin content originating from the flow stagnation that may provide clot structural support at the base of the injury site. In reality, we anticipate that the pooled flow and fibrin formation processes will also extend into the extra-vascular space from the injury site. In prior work, we have computationally demonstrated potential mechanisms by which the flow leakage from the base of the clot may influence clot flow interactions [29]. While we have not focused on illustrating this phenomena in detail in this specific study, there are simple avenues to incorporate estimates of leakage flow from image processing and segmentation of the cAlb concentrations. Further characterization of extended shell and extra-vascular space can be reasonably undertaken through numerical experimentation using additional sample mouse datasets based on the methodology developed here.

**Figure 8:**
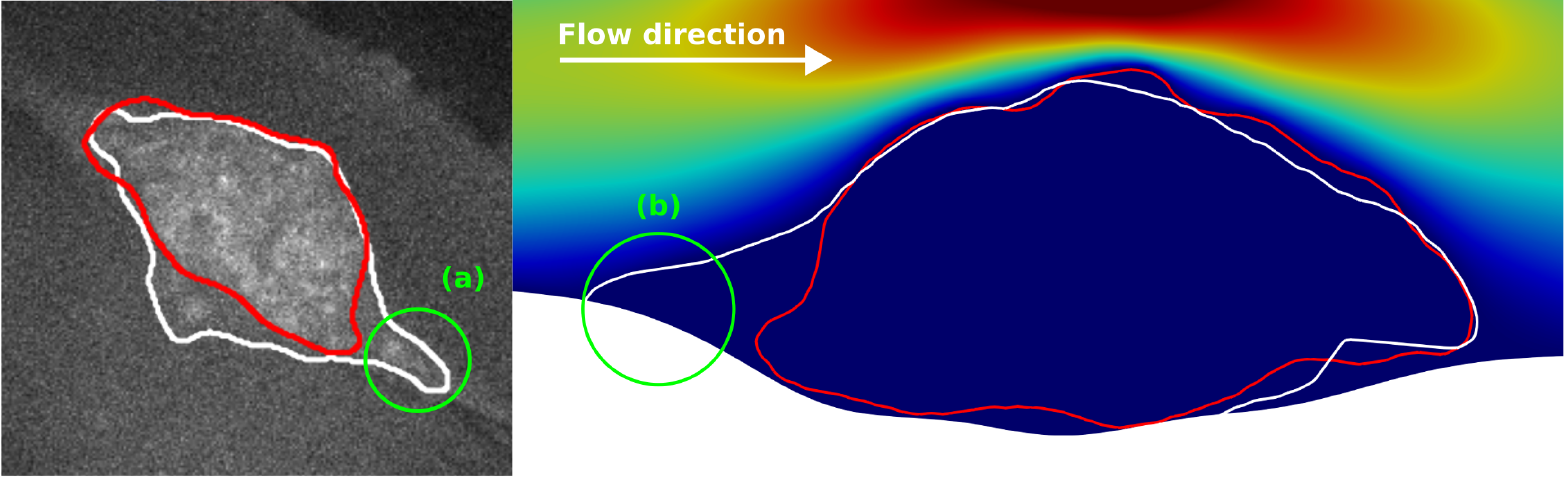
Sample illustration of the slow arrested flow region at the base of the clot referred to as the extended shell (labelled as (a) and (b) on the two figure panels). On the left panel, (a) labels the segmented outline of high intensity of cAlb signal, which is outside the clot shell domain (marked in red segment), and which is identified with slow stagnated flow on the right panel labeled by (b).

The analysis of flow-induced clot deformation properties as presented in Section 3.4 and Figure 7 enables exploring a key question regarding the extent of force that the clot can sustain prior to significant stretching, deformation, and embolization. In our `IVISim` simulation case studies we do not require a clot material model as we reconstruct clot kinematics and deformation at a high fidelity directly from images. However, we do characterize the clot heterogeneous composition using microstructural parameters like porosity, and core-shell variations. For a given microstructure, a stable clot can be expected to withstand a greater extent of flow-induced loading as compared to a structurally weaker clot. Comparisons between WT and diYF knockout models provide some evidence of this behavior as shown in Figure 7. This is of major significance, indicating that `IVISim` can potentially enable investigations on arterial clot stability *in vivo*, which remains currently a major challenge as indicated in several works [5, 51]. We also note that `IVISim` is proposed as a methodology to evaluate the flow and transport environment for known clot growth and deformation kinematics as estimated directly from intravital images, leveraged the clot structure information to bypass requiring a material model. Consequently, this is demonstrated here as a quasi one-way fluid-structure interaction approach, where we do not explicitly model the deformation mechanics using clot material properties. Hence, we have not addressed factors like determination of clot elasticity modulus, yield stress, and viscous behavior. While this is a limitation of the analysis presented here, we can use `IVISim` to set up systematic inverse problems for identifying clot material models and parameters. Specifically, an avenue for future investigation could involve inverse investigations of choice of material models or material parameters that correspond to a known clot domain dynamics for a known state of flow-induced stresses on the clot. Such investigations will require significant theoretical and computational effort which render them beyond the scope of this current study.

The clot porosity and diffusivity values used here were estimated based on prior literature. We note that in reality for a dynamically growing, deforming, and embolizing clot the porosity and diffusivity can both vary spatio-temporally, implying that *ϕ* = *ϕ* (***<u>x</u>***, *t*) and *D* = *D* (***<u>x</u>***, *t*) for *in vivo* clots as outlined in Section 2.5. These spatio-temporal microstructural variations are governed by intricate interplay of clot deformation dynamics, physiological clot consolidation, and a slew of biochemical and genetic factors. Here, we have considered variabilities in terms of core and shell microstructure as outlined in 2.5.2, but no added variabilities based on detailed pore network architecture and other structural and compositional features. This is a limitation of the current study. However, this is not an inherent limitation of the `IVISim` framework as developed here. When such detailed spatio-temporally resolved characterizations of the microstructure are available, the `IVISim` framework can incorporate this information owing to the custom stabilized finite element algorithms implemented within the framework. However, such dynamic high-resolution space-time varying characterization of the clot microstructure *in vivo* is challenging to obtain. Hence, in this study we have demonstrated our simulation cases by resolving the core and shell structure as a starting point. In the absence of such experimental data, brute force simulation efforts for statistical characterization and uncertainty quantification for these spatio-temporally varying microstructures will require a significantly comprehensive computer simulation effort spanning a range of representations of microstructure and diffusivity, and over multiple mouse or microfluidic models. While this was beyond the scope of the present study, the aspects outlined here remain a major motivation for future investigations based on `IVISim` capabilities. On a related note, in prior works [28], we have shown that for small variations in microstructural features, the flow patterns and resultant pressure loading remains relatively unaffected. However, large spatio-temporal variabilities in clot porosity and other microstructural parameters may lead to variabilities in the estimated flow-induced loading; which requires systematic sensitivity analysis. While a comprehensive sensitivity analysis spanning the entire set of parameters from image-processing to transport variables is a commensurately complicated endeavor, we have demonstrated here the sensitivity analysis for the image-processing modules of the framework via integration with a specialized library `Dakota`. This can further be exploited with techniques of surrogate modeling and reduced order computations to set up efficient sensitivity and uncertainty quantification in future efforts.

The capability for dynamic reconstruction of coupled flow, transport, and clot deformation can enable key advancements in hemostasis and thrombosis with clinical implications. This can involve mechanisms and etiology of related pathological manifestations, as well as efficacy, outcomes, and risks associated with pharmacological or endovascular interventions [2]. For example, the elution and local transport of thrombin *in vivo* is central to hemostatic response [52]. Advancements in understanding of thrombin transport and distribution, as well as local transport of key biochemical agonists such as ADP and Thromboxane can drive advancements in approaches for hemostatic therapy [53, 54]. As another example, advancements in understanding of structure and stability of a thrombus or thrombo-embolus can have major clinical implications in terms of endovascular thrombectomy procedures and distal embolization risks in stroke [55]. Additionally, the interplay of transport with clot microstructural composition is increasingly recognized to be highly relevant for thrombolytic therapy [56] - in terms of understanding how perfusion of tPA and subsequent fibrinolysis occurs, as well as in terms of understanding how novel targets such as Von Willebrand Factor (VWF) can be accessed for pharmaceutical interventions [7] *In silico* integration and supplementation of *in vivo* and *in vitro* data as facilitated by `IVISim` framework can transform into a significant modality for further investigations focused on such aspects of clinical and therapeutic relevance.

## Supporting information

Supplemental Animation: sup-movie-diyf

Supplemental Animation: sup-movie-wt

## Acknowledgments

This work utilized resources from the University of Colorado Boulder Research Computing Group, which is supported by the National Science Foundation (Awards: ACI-1532235 and ACI-1532236), the University of Colorado Boulder, and the Colorado State University. Collaborative discussions between Authors DM, MT, and TS were partly initiated and supported by a Burroughs Wellcome Fund Collaborative Research Travel Grant (Award: 1016360), and an American Heart Association award (Award: 16POST27500023) - both with DM as principal investigator.

## Author contributions

CT and DM developed the image-processing and finite element algorithms. CT implemented the complete workflow, conducted all simulations and data analysis. MT and TS identified intravital experiment cases, post-processed intravital data, and guided intravital to in silico translation. CT and DM developed the manuscript with technical inputs and insights from MT and TS. All Authors reviewed the manuscript and agreed to the final version as submitted.

## Data availability

All simulation data generated or analyzed during this study are included in this published article and its supplementary information files. The simulation codes generated and/or analyzed during the current study are not publicly available due to the reason stated as follows, but are available from the corresponding author on reasonable request. The underlying source codes for the numerical framework reported here are currently being packaged and built into a software library for open source release. Up until that time, research purpose access to the development version of the source codes can be obtained by directly emailing the corresponding author or by sending a message through the following url: https://www.flowphysicslab.com/contact.html.

## Supplementary Material Information

### S1 Image pre-processing algorithm implementation

Since the pixel intensity value of the noise is comparable to those inside the clot domain, running a general-purpose denoising algorithm on the image directly does not remove the noise effectively. To alleviate this problem, we binarize the image such that the pixel intensity above a specific intensity value is set to 1 and the rest is 0. In this work, we binarize the image based on the pixel intensity distribution of the image according to the following formula

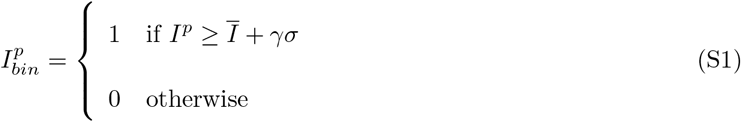

Where *I*^*p*^ is the intensity of pixel *p*, 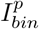 is the binarized version of pixel *p*, Ī is the mean pixel intensity value of the image, *σ* is the standard deviation of the image’s pixel intensity distribution, and *γ* is a constant parameter that can be tuned. A typical *γ* value used in this study is 1.5. As shown in panel *b* of figure S1, binarizing the image differentiates the bright, sparsely spaced (noisy) regions from the bright, tightly packed thrombus region.

### S2 Automated clot boundary selection algorithm

After the image frames are pre-processed, and the boundary is segmented, the ending result may contain multiple disconnected segments (see panel c of figure S2). In this work, we pick the boundary with the largest perimeter as the final clot domain boundary. To perform this selection automatically, the boundary pixels are represented as a graph where each boundary pixels are the graph vertex. Vertex connectivity is defined as pixels that share a corner or an edge. The algorithm performs a depth first search (DFS) graph traversal starting from an initial vertex *v*_0_ until every vertex in the locally connected graph is visited. These visited vertices are marked with an index value, and a new starting vertex *v*_0_ is selected from the list of un-visited vertices. The algorithm continues until every boundary pixel on the graph is visited. The boundary with the largest number of boundary pixels is selected as the clot boundary.

### S3 Clot boundary interpolation between time steps

Points along the boundary are linearly interpolated in time to obtain the clot boundary geometry between subsequent time increments. Since the clot boundary is a closed curve for our model purposes, we interpolate between points on the boundary with the same angle to the geometric center of mass of the clot. This process can be seen as drawing rays radially outward from the clot geometric center of mass and interpolate clot boundary points between intersecting points along those rays S3. A tree search algorithm is used to efficiently locate the intersection point between each ray and the B-Spline curve that represents the boundary of the clot. The tree search recursively performs local curve refinement until an intersection point with the desired tolerance is found. The complete algorithm description for the tree search is provided in Algorithm 2. The interpolation accuracy of the clot boundary across time increments was further examined by down sampling the image data and re-interpolating the temporally down sampled images between each frame. The data was down sampled by removing every two frames between the images. The boundary of the removed frames were then interpolated using the `IVISim` framework, and examined against the original boundary as directly obtained from the original image stack. The down sampling analysis and comparison was performed on the CD41 channel as the shell of the clot has the most dynamic features. Figures S4 and S5 present the interpolated boundary from the down sampled image (*marked in yellow lines*), and the extracted boundary from the original image stack (*marked in red line*), for both WT and diYF knockout cases respectively. We observe that the interpolated boundaries show excellent agreement.

### S4 Details of finite element numerical formulation

#### S4.1 Blood flow numerical formulation

Blood is assumed to be an incompressible, Newtonian fluid. Flow around the thrombus is computed using the unsteady Navier-Stokes equation. Flow inside the porous thrombus domain is modeled using the Brinkman equation for low Reynolds number flow inside porous media [39]. The system is solved using a Petrov-Galerkin stabilized finite element. Briefly, let Ω be the fluid (blood) domain, and Γ = *∂*Ω is the domain boundary. The flow equations are solved using a Petrov-Galerkin stabilized formulation of the Navier-Stokes equation with the overall weak form stated as follows:

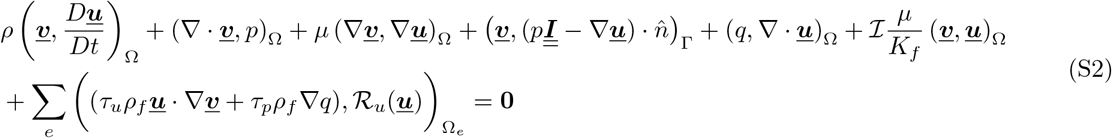

where we use the notation (*a, b*)_Ω_ = ∫ _Ω_ *a b d*Ω; *μ, ρ* are averaged whole blood dynamic viscosity and density respectively; ***<u>u</u>***, *p* are the velocity and pressure trial functions; ***<u>v</u>***, *q* are the respective test functions; ℛ_*u*_ is the momentum residual; *τ*_*u*_ and *τ*_*p*_ are the velocity and pressure stabilization terms [40]; *K*_*f*_ is thrombus permeability. Thrombus permeability is modeled as a function of thrombus porosity *ϕ* using the Kozeny-Carman relation:

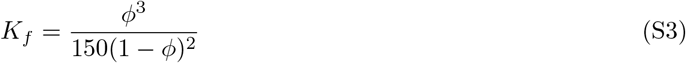

#### S4.2 Chemical transport numerical formulation

The mass transport problem is solved using the SUPG stabilized unsteady advection diffusion equation. We model hindered diffusion inside the clot domain by assigning different diffusivity value inside the clot. Discontinuity in the diffusivity field leads to instability which requires additional stabilization. In this work, we use the continuous interior penalty stabilization method outlined in [41] which are briefly defined as follows

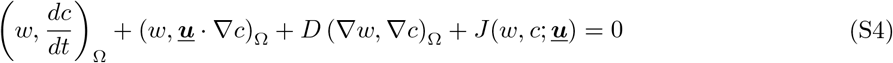

where *w* and *c* are the standard test and trial functions of the concentration field respectively; ***u*** is the background flow velocity; *J*(*w, c*) is the continuous interior penalty stabilization parameter defined as

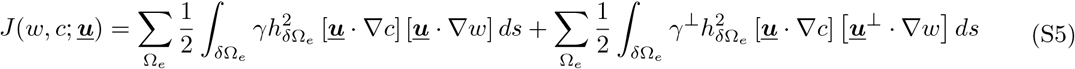

where 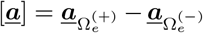 is the jump of value ***<u>a</u>*** across element facet 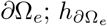 is the measure of facet *∂*Ω_*e*_; variable ***<u>u</u>***^⊥^ is a vector perpendicular to the flow velocity ***<u>u</u>***; *γ* and *γ*^⊥^ are constant parameters.

### S5 Additional illustrations

Here we include two additional illustrations that supplement the data and content in the main manuscript. First, Figure S6 supplements the observations discussed in Section 3.4 and Figure 7. Here we provide estimates of the pressure force, the shear force, and the sum total of both for the WT and diYF mouse clots studied in this manuscript. The panels a-c in Figure S6 establishes that the pressure force is dominant compared to the shear force, and justifies our choice of including the total hemodynamic force to account for relative variabilities between pressure and shear. Second, Figure S7 supplements the observations presented in Section 3.1 regarding comparisons of trends in clot kinematics as extracted using the automated approach in `IVISim` and those obtained from averaging of clot kinematics extracted using a non-automated method across multiple mouse experiments.

### S6 Sensitivity analysis for image processing

In this analysis, we examine the sensitivity of the segmented image area and aspect ratio to the image processing parameters for `IVISim`. For this purpose, we implemented an integration of the `IVISim` codebase with the open source library `Dakota` [47, **?**]. `Dakota` is a specialized library for conducting parameter sensitivity, uncertainty quantification, and optimization. For our sensitivity analysis, the sampling of image processing parameters was done using the Latin hypercube sampling technique implemented within `Dakota`. The bounds of the image processing parameters distributions were chosen based on an initial manual inspection of the segmentation results to ensure no unfeasible parameter ranges are specified as inputs.

#### S6.1 Analysis with synthetic image data

To examine the sensitivity to image processing parameters, we developed a set of synthetic ellipsoidal clot image data with added noise. The added noise over the entire analysis domain is a Gaussian noise centered at 0 with standard deviation equal to the standard deviation of the microscopy image pixel intensities. The background intensity level of the overall image is the mean intensity value of the microscopy image pixels outside of the imaged clot domain, and the ellipse pixel intensity is two standard deviations higher than the background intensity. With the synthetic ellipsoidal image based tests, we have the advantage of knowing an exact clot area, aspect ratio, and boundary locations, which can be compared against the segmented value using our methodology. For our analysis, we chose synthetic elliposidal images with aspect ratio equals to 0.5, 0.75, and 1.0, which correspond to Figures S8, S9, and S10 respectively. The panel b. represents the ellipse geometry on a background grid, the panel c. represents the synthetic image version of the ellipse that is used for sensitivity analysis. Panel a. in these figures represent the output from the sensitivity analysis conducted through the integration of `IVISim` with `Dakota`. The analysis tested variations in the four key parameters for the image processing module as described in Section 2.2, S1, and S2: namely threshold *γ*, median filter size, Gauss filter size (or blur size), and Gauss filter standard deviation (or blur strength). We quantified area and aspect ratio of the segmented image with reference to that of the exact specified ellipsoid, and in general observe that the segmentations reproduce the same ellipsoid with an error range of 5-10% as observed across Figures S8, S9, and S10.

#### S6.2 Analysis with microscopy image data

We further extended the sensitivity analysis to image processing of the microscopy data. The same procedure stated above has been followed, but for images at individual instances in time. Figure S11 presents the results from the `Dakota` Latin Hypercube Sampling analysis for one such sample image at one instant in time. Panel b. shows the overlay of multiple segmentation outputs all superposed into one image, and we observe that the segmented outline (marked in red) is very robust across changes to the four image processing parameters identified above. For this image, the main variations are observed in the low intensity tail/distal region of the clot. Panel a. shows that within the chosen range of image processing parameters, the resulting area varies within approximately ≤ 10% of the mean value, and aspect ratio varies within approximately ≤ 5% of the mean value. Here, we note that the computed aspect ratio is the aspect ratio of the segmented boundaries oriented as shown in Figure S11b.

### S7 Additional illustrations: Animations

Two different animations have been provided as supplementary information for this manuscript, to visualize the unsteady clot hemodynamics interactions obtained from `IVISim` workflow. The first animation movie file named: ***sup-movie-wt*.*mp4*** demonstrates the evolution of clot kinematic and force-deformation through time for the wild type (WT) mouse model described in the manuscript. The second animation movie file named: ***sup-movie-diyf*.*mp4*** demonstrate the same for the diYF knockout mouse model described in the manuscript. The clot image presented is that of the CD41 channel which constitute the solid region of the clot domain. Each animation provides also the dynamic computed predictions of clot shape and flow-induced stresses on the clot boundary, for a total duration of 165 seconds (that is 2 mins and 45 seconds) of real time.

### S8 List Of Algorithms

#### Algorithm 1

Algorithm implemented for the selection of largest boundary segment in the `IVISim` automated image processing module.

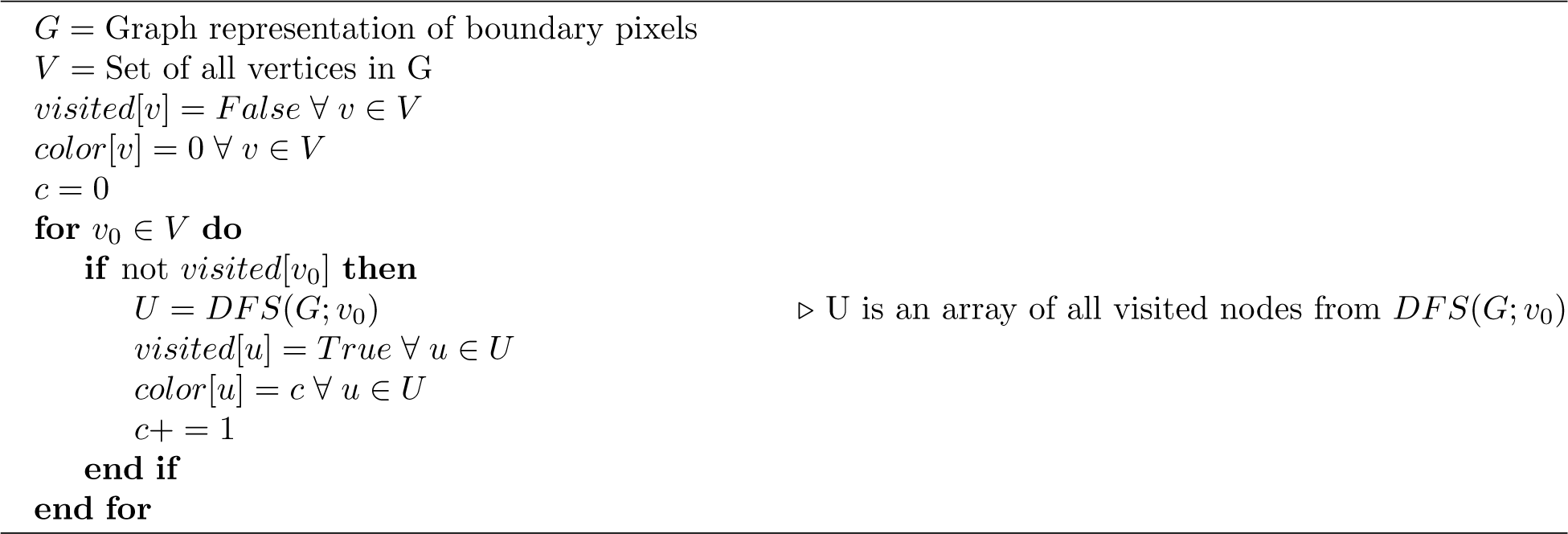

#### Algorithm 2

Algorithm implemented for B-spline - ray intersection within `IVISim` automated image processing module

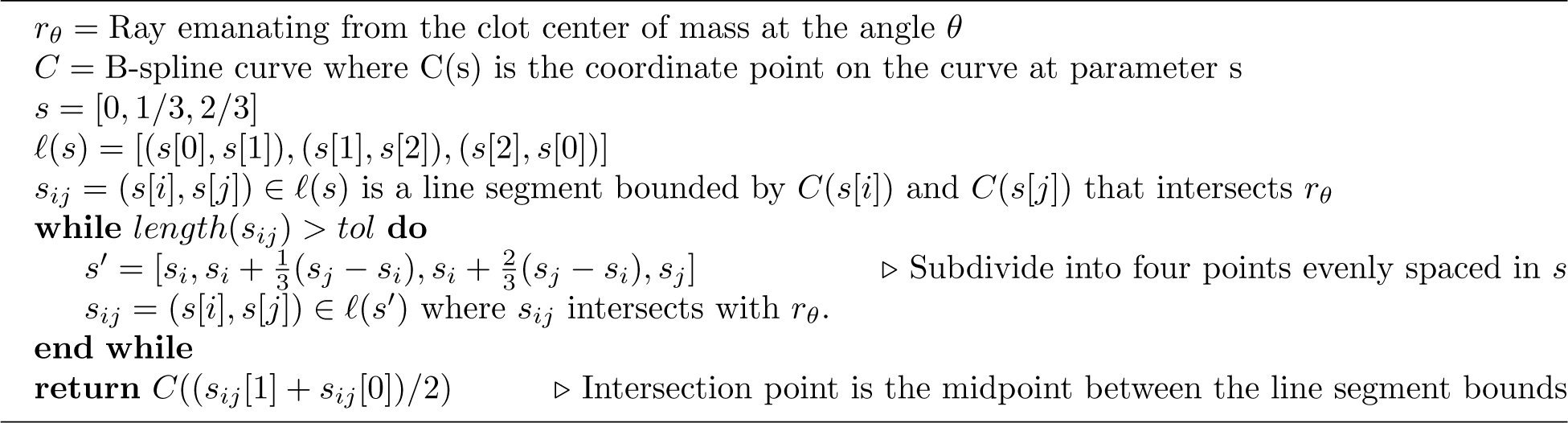

### S9 All Supplementary Figures

**Figure S1:**
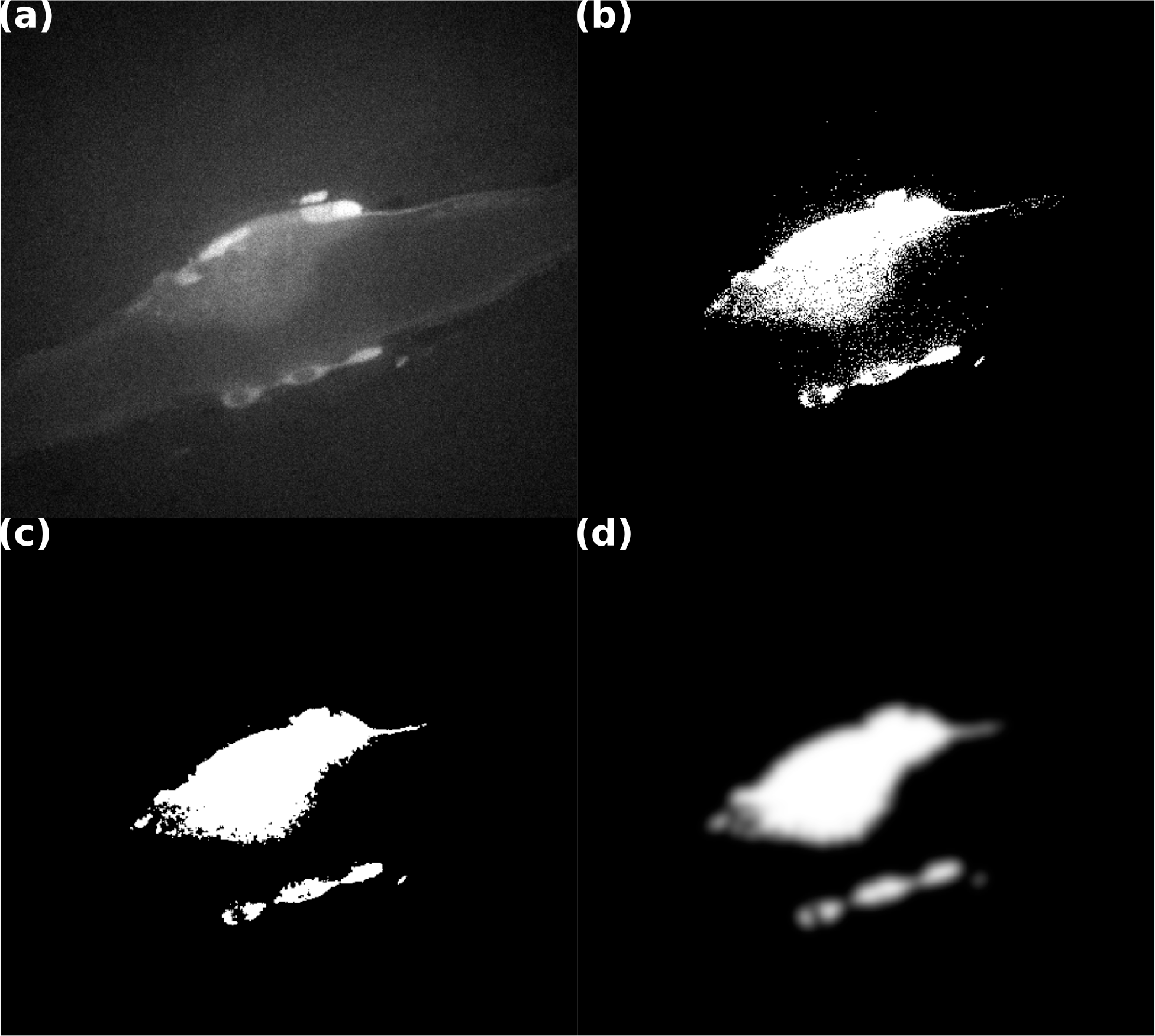
(a) Raw image is obtained from intravital microscopy experiment; (b) image is binarized to remove most of the non-clot domains; (c) salt and pepper noise is removed using median filter; (4) clot is smoothed out using gaussian filter.

**Figure S2:**
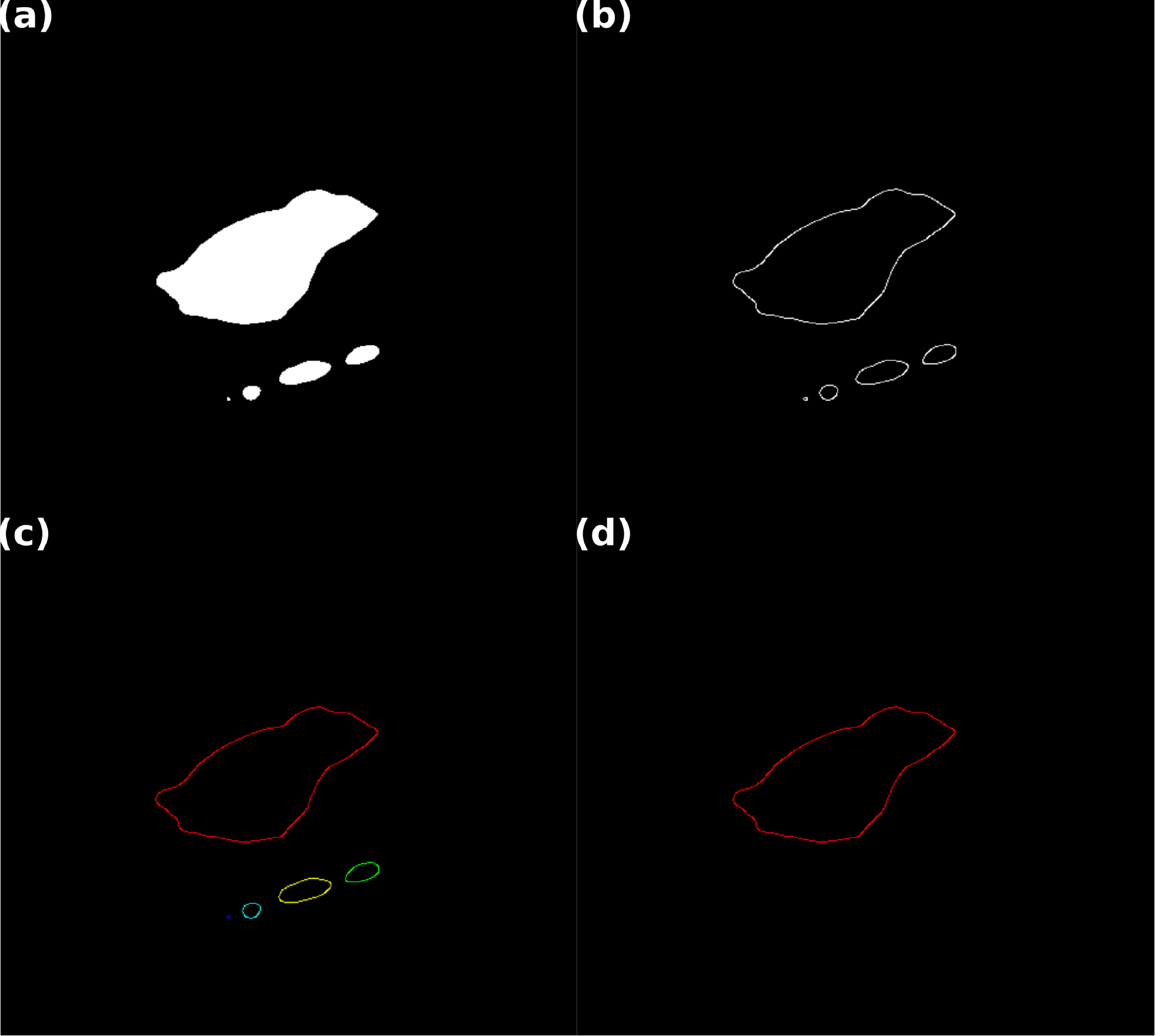
(a) Filtered image is segmented using k-means clustering; (b) canny edge detection is used to get boundary pixels; (c) disconnected boundary are sorted by its perimeter length; (d) the boundary with the largest perimeter is selected as the clot domain.

**Figure S3:**
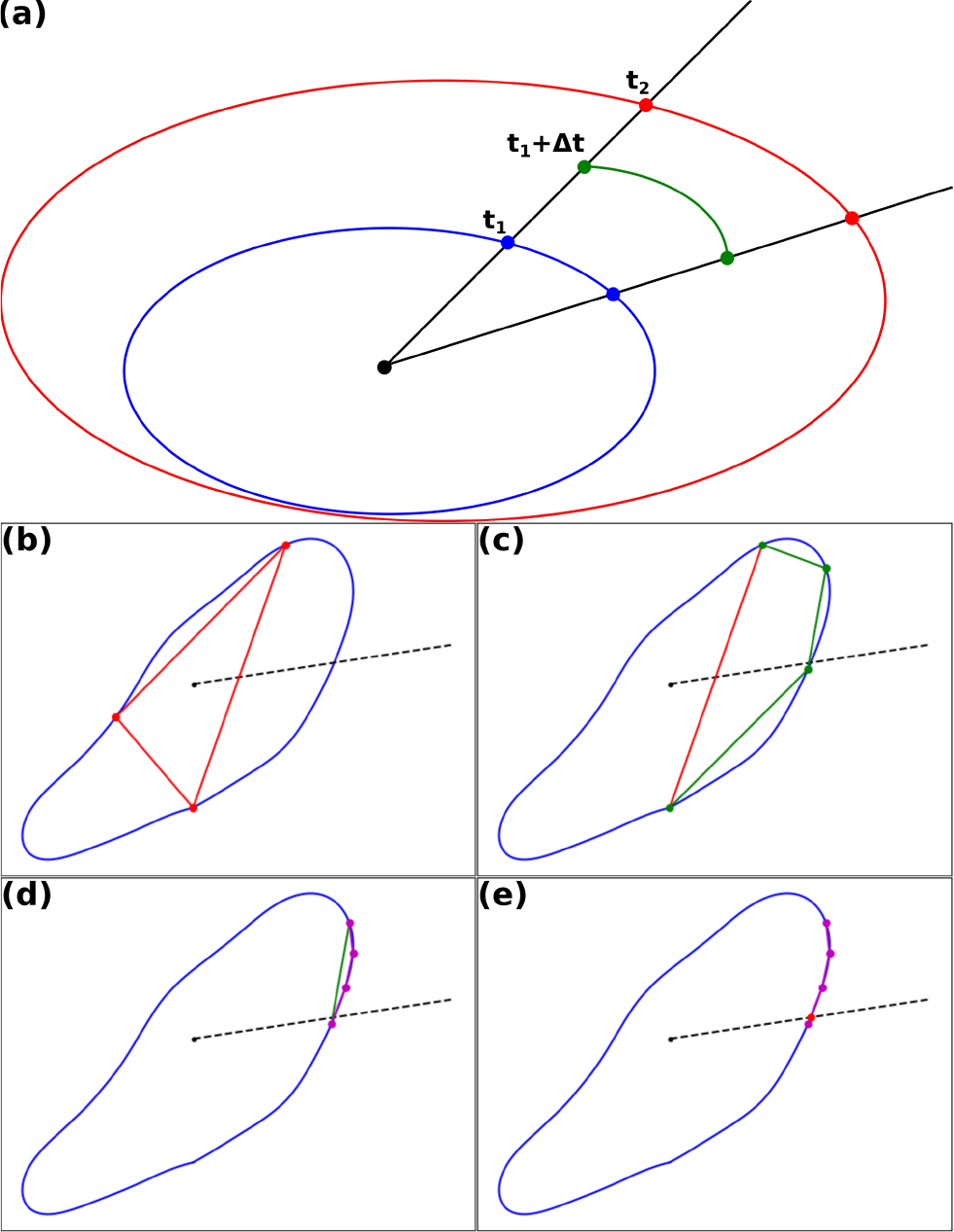
(a) Clot boundary is interpolated between frames by interpolating between points intersecting a ray emanating from clot center; (b,c,d,e) intersection point is efficiently computed by successively reducing line segment size.

**Figure S4:**
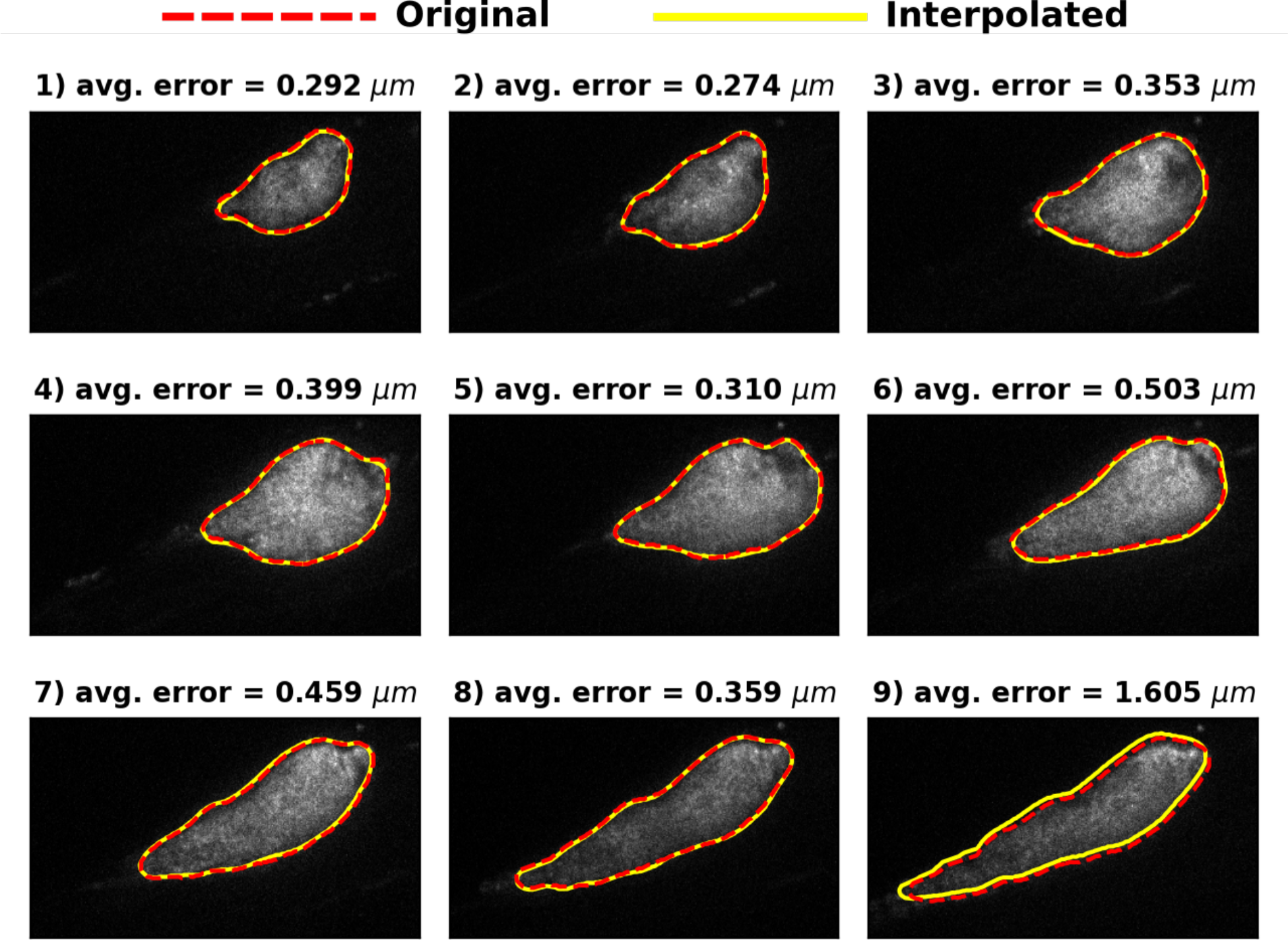
Interpolated boundary from a down sampled image frames compared to the original segmented boundary data for the wild type data set. The avg. error is the average point-wise distance from the original and interpolated boundary along the same interpolating ray.

**Figure S5:**
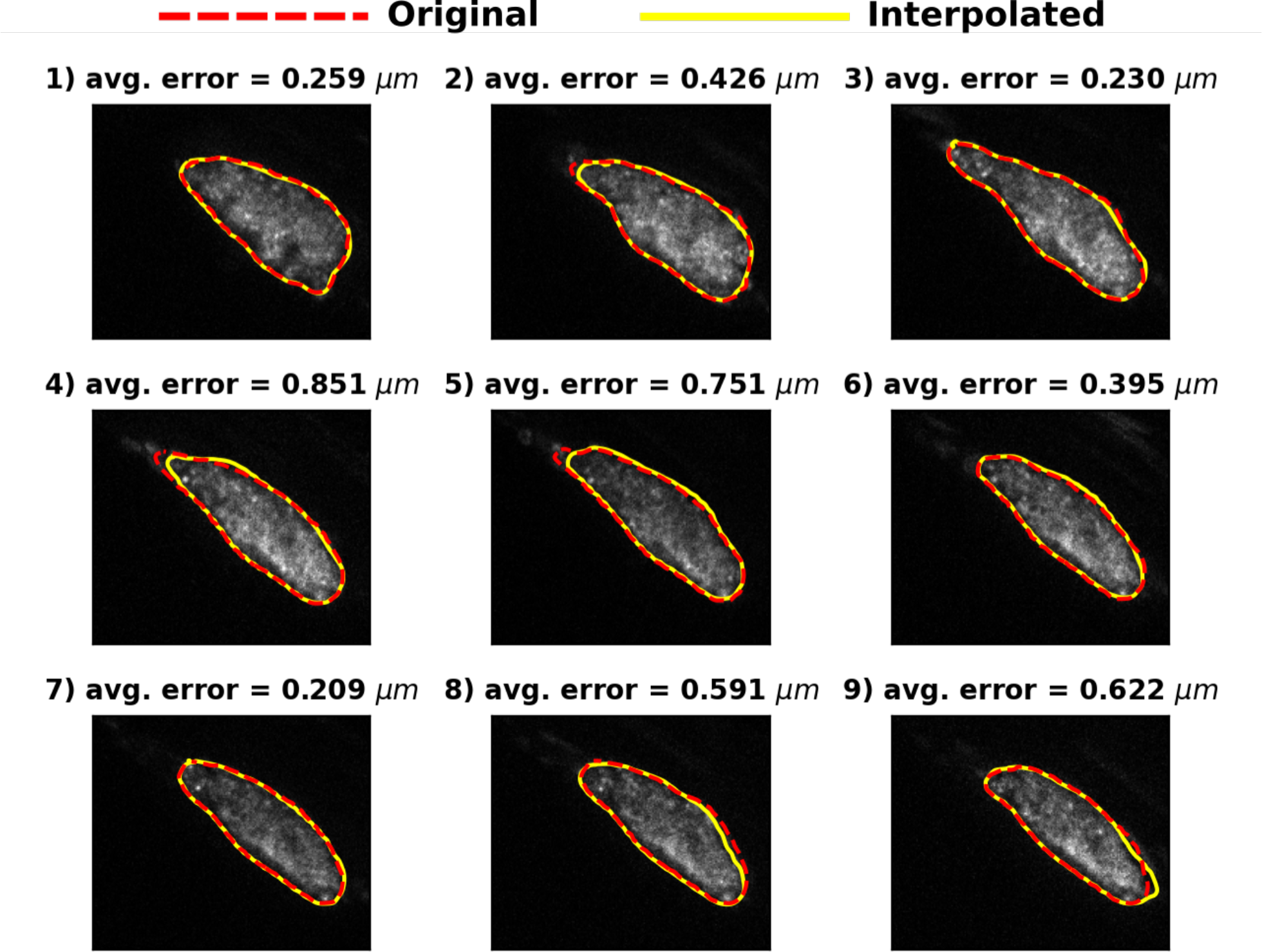
Interpolated boundary from a down sampled image frames compared to the original segmented boundary data for the diYF data set. The avg. error is the average point-wise distance from the original and interpolated boundary along the same interpolating ray.

**Figure S6:**
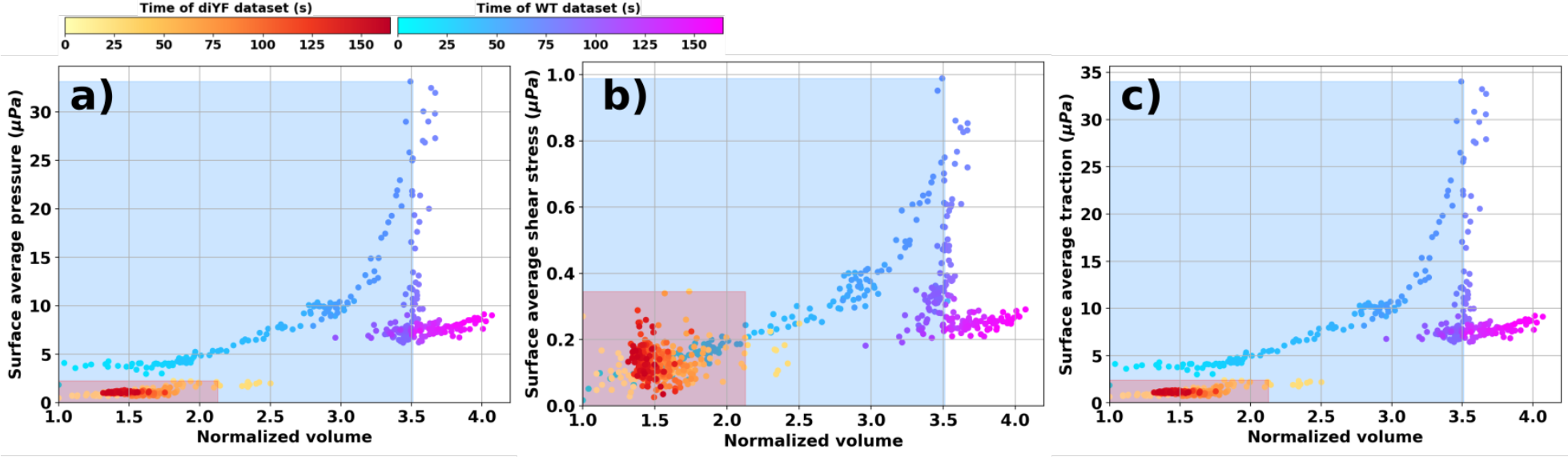
Components of the clot-hemodynamic interaction force and clot force-deformation behavior: a) pressure force; b) shear force; c) total force.

**Figure S7:**
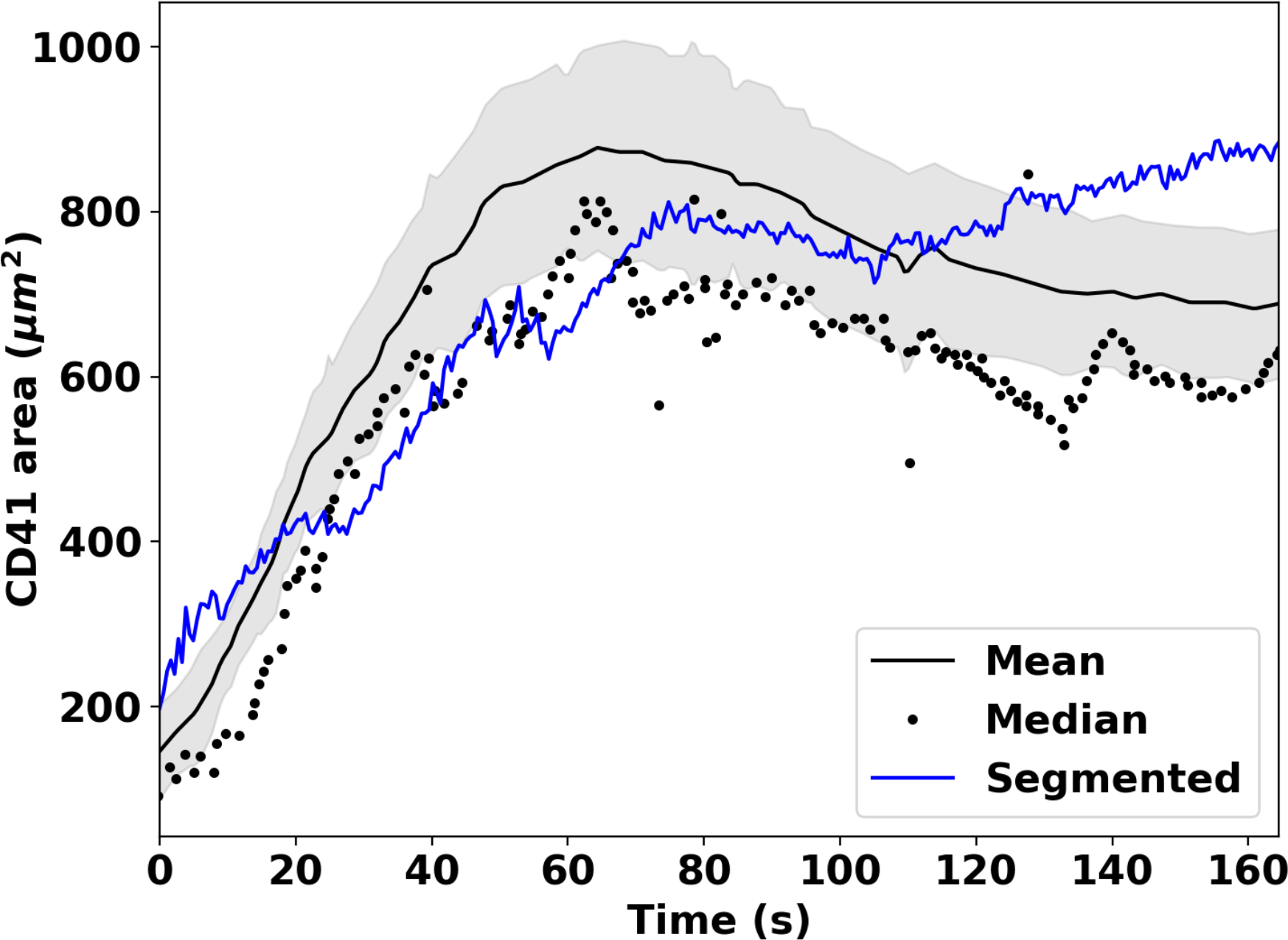
Illustration of a quantitative comparison for the one WT mouse model sample clot domain kinematics data against the averaged clot domain kinematics trends as reported in [30], where the clot domain is interpreted from the CD41 fluorescence signal intensities. The more accurate automated reconstruction approach outputs agree reasonably well with the statistically averaged trends from multiple mice, obtained using a non-automatic reconstruction algorithm.

**Figure S8:**
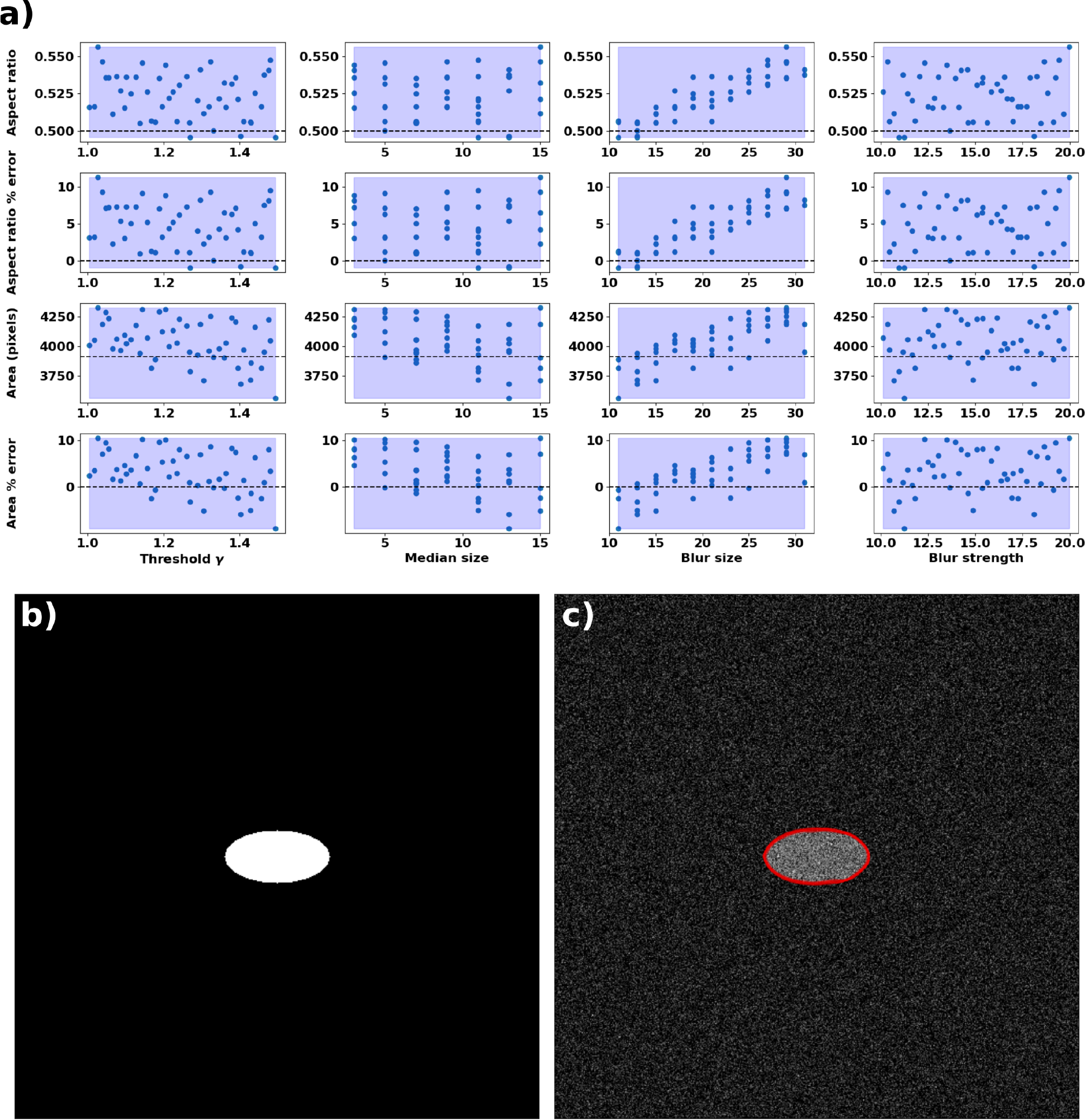
a) Sensitivity of the segmented boundary; b) baseline synthetic image; c) noisy image with segmented boundary overlay.

**Figure S9:**
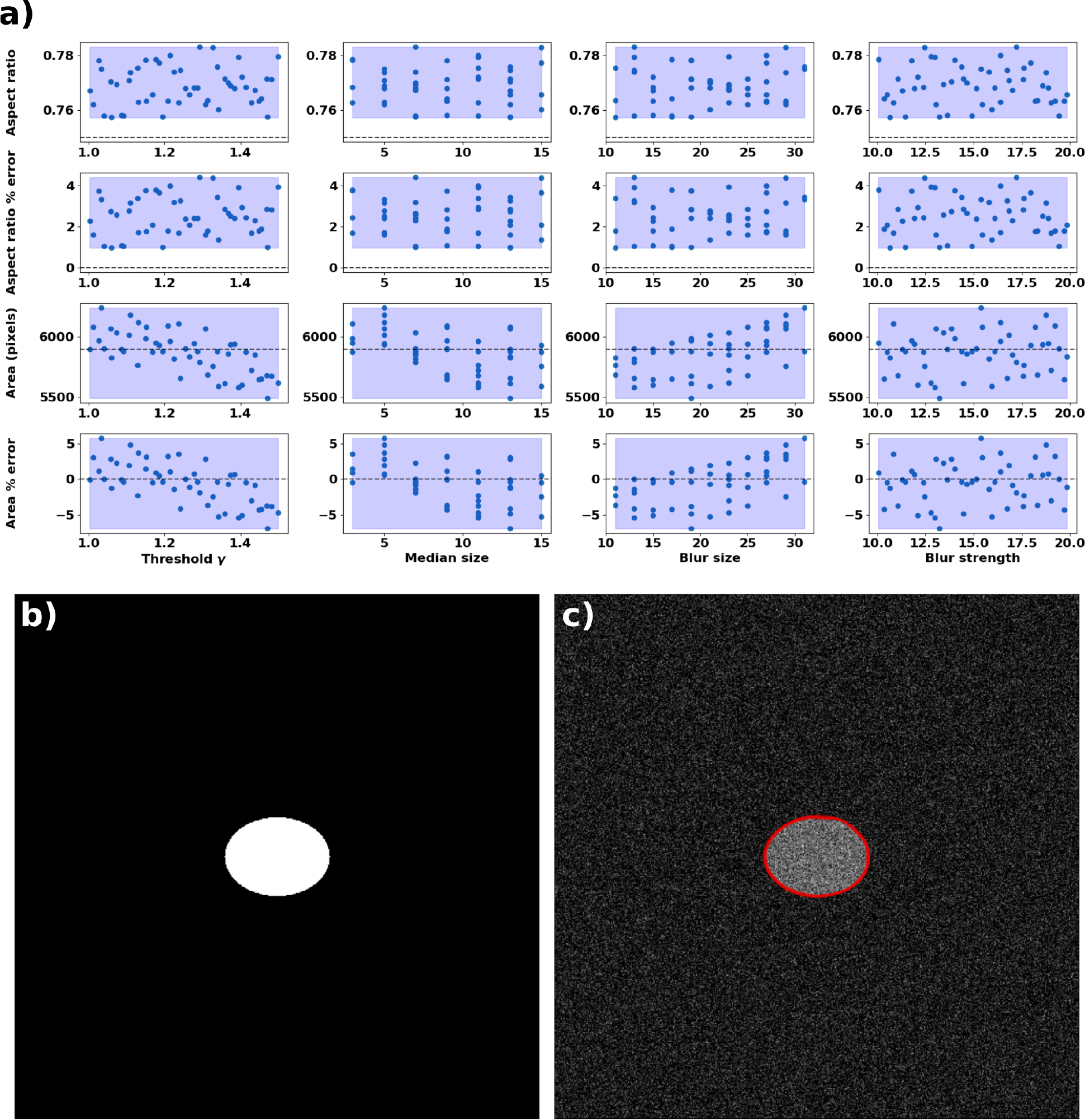
a) Sensitivity of the segmented boundary; b) baseline synthetic image; c) noisy image with segmented boundary overlay.

**Figure S10:**
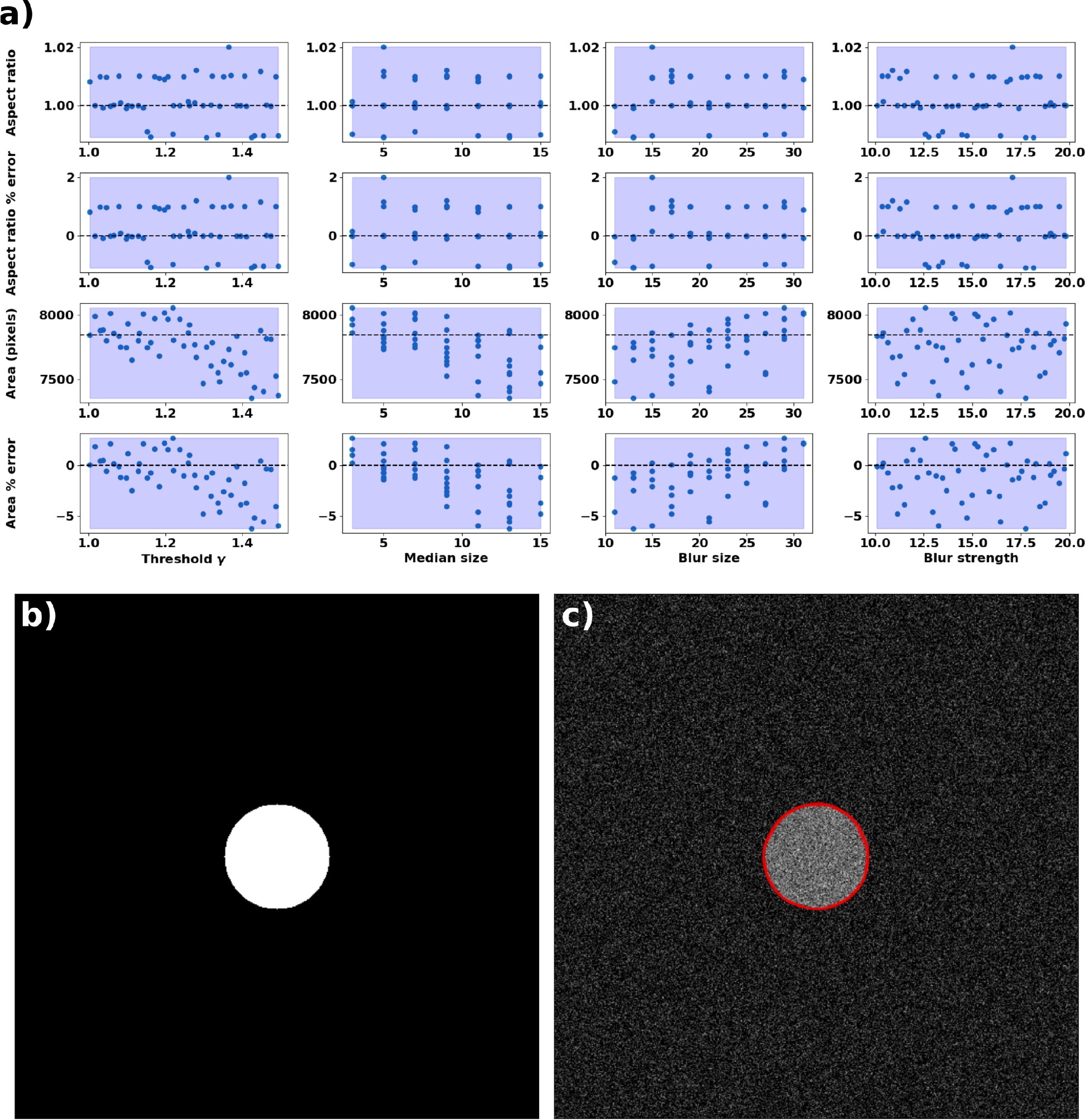
a) Sensitivity of the segmented boundary; b) baseline synthetic image; c) noisy image with segmented boundary overlay.

**Figure S11:**
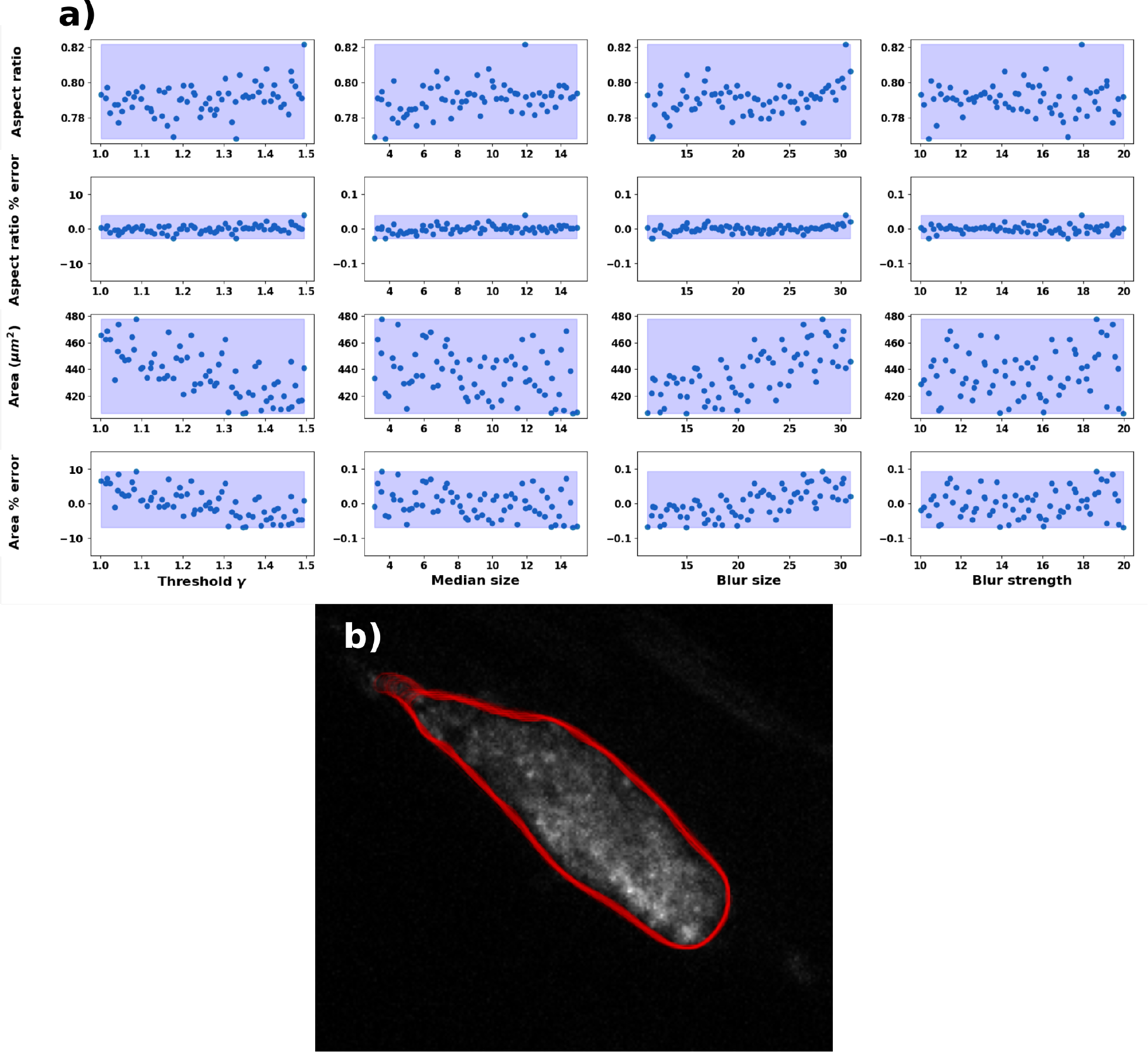
CD41 channel a) Variability of the segmented boundary based on varying image segmentation parameters; b) all segmented boundaries overlay on the clot image.

**Figure S12:**
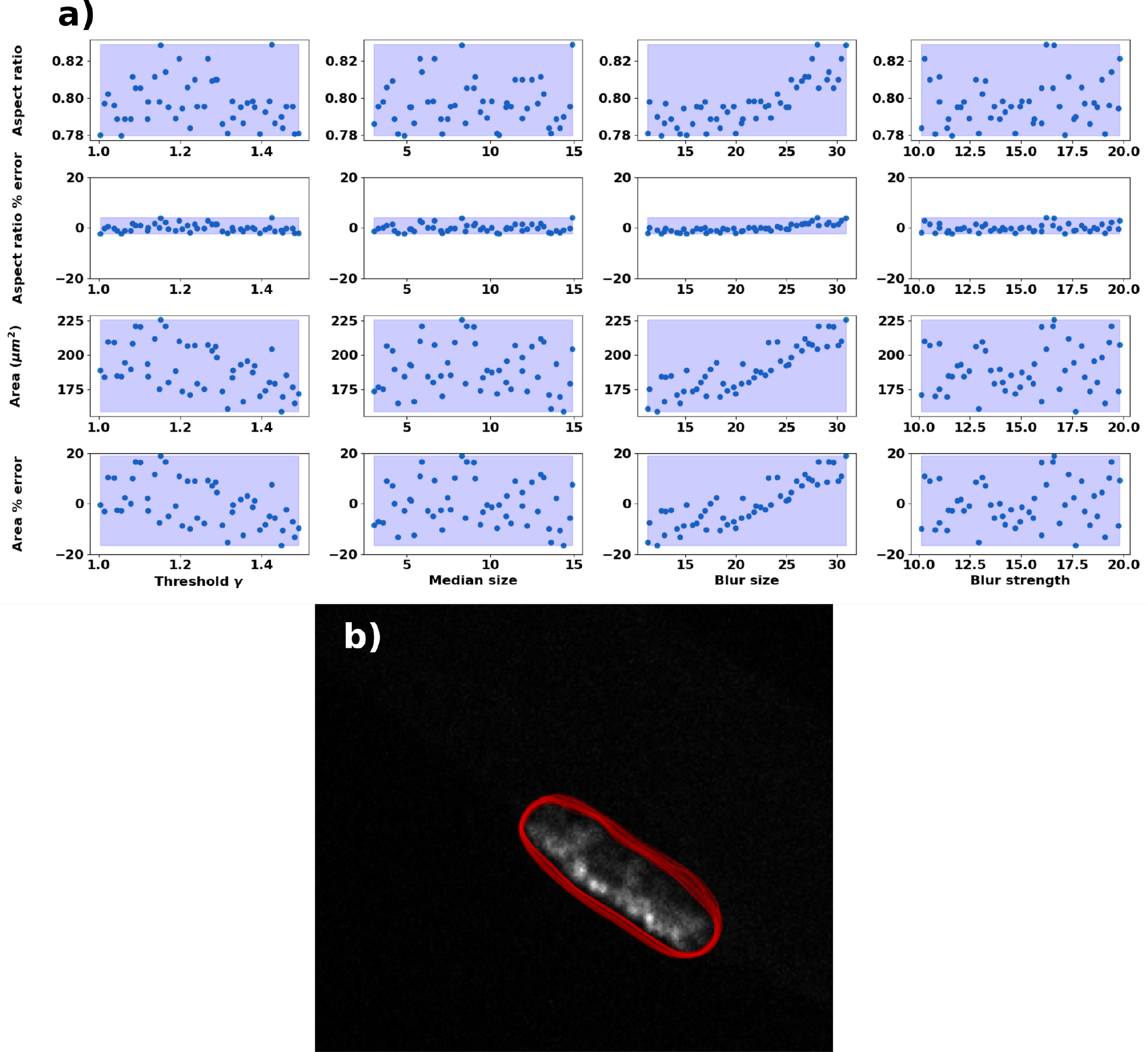
P-Selectin channel a) Variability of the segmented boundary based on varying image segmentation parameters; b) all segmented boundaries overlay on the clot image.

